# Unlocking genome-based prediction and selection in conifers: the key role of within-family prediction accuracy

**DOI:** 10.1101/2024.01.25.577259

**Authors:** Victor Papin, Gregor Gorjanc, Ivan Pocrnic, Laurent Bouffier, Leopoldo Sanchez

**Affiliations:** INRAE, BIOGECO, UMR 1202, 69 route d’Arcachon, 33610 Cestas, France. University of Bordeaux, BIOGECO, UMR 1202, 33400 Talence, France; The Roslin Institute and Royal (Dick) School of Veterinary Medicine, The University of Edinburgh, Edinburgh, UK; INRAE-ONF, BioForA, UMR 0588, 2163 Avenue de la Pomme de Pin, CS 40001 Ardon, 45075, Cedex 2, Orléans, France

**Keywords:** Breeding programme, genomic selection, maritime pine, progeny validation, stochastic simulation, within-family variability

## Abstract

**Context:** Genomic selection is a promising approach for forest tree breeding. However, its advantage in terms of prediction accuracy over conventional pedigree-based methods is unclear and within-family accuracy is rarely assessed.

**Aims:** We used an pedigree-based model (ABLUP) with corrected pedigree data as a baseline reference for assessing the prediction accuracy of genome-based model (GBLUP) at the global and within-family levels in maritime pine (*Pinus pinaster* Ait).

**Methods:** We sampled 39 full-sib families, each comprising 10 to 40 individuals, to constitute an experimental population of 833 individuals. A stochastic simulation model was also developed to explore other scenarios of heritability, training set size and tagging density.

**Results:** Prediction accuracies with GBLUP and ABLUP were similar and accuracy with GBLUP within-family was on average zero with large variation between families. Simulations revealed that the number of individuals in the training set was the principal factor limiting GBLUP accuracy in our study and likely in many forest tree breeding programmes. Accurate within-family prediction is possible if 40-65 individuals per full-sib family are included in the genomic training set, from a total of 1600-2000 individuals in the training set.

**Conclusion:** Such conditions lead to a significant advantage of GBLUP over ABLUP in terms of prediction accuracy and more clearly justify the switch to genome-based prediction and selection in forest trees.

## 1. Introduction

The use of pedigree information in populations with genealogy records has improved breeding programmes for many species. Pedigree information can be used to infer the expected relatedness between each pair of individuals, making it possible to gauge the extent to which phenotypic values of individuals in the studied population have a genetic basis. This pedigree-based model underpinned the development of predictions of the individual additive genetic value, or the breeding value, via BLUP (Best Linear Unbiased Prediction) methodology (Henderson, 1975). Breeding value of each individual can be decomposed into parent average and Mendelian sampling terms. Parent average term captures variation between families, it represents expected breeding value of progeny given its parents. Mendelian sampling term captures variation within families, it represents deviation of each individual’s breeding value from the parent average due to recombination and segregation of parental genomes. With pedigree-based model, we need phenotypic values on an individual or its progeny to estimate the Mendelian sampling term of the individual’s breeding value. Hence, pedigree-based prediction of breeding values for non-phenotyped individuals (forward prediction) captures only parent average term. By using genome data, we observe outcome of recombination and segregation of parental genomes as well as recent or past mutations, meaning that we can in principle estimate parent average and Mendelian sampling terms of breeding value even for non-phenotyped individuals. (VanRaden, 2008, Hill & Weir, 2011).

The advent of affordable genome-wide DNA marker genotyping platforms has enabled such genome-based predictions paving the way to genomic selection (GS). Albeit initially proposed by Bernardo (1994) and Nejati-Javaremi et al. (1997), genome-based prediction took off with the work of Meuwissen et al. (2001), which showed how regressing individuals’ phenotypic values onto their genome-wide marker genotypes captured variation between individuals’ breeding values by leveraging linkage-disequilibrium between quantitative trait loci (QTL) affecting traits of interest and the genome-wide markers. Using a training set of individuals that have been phenotyped and genotyped, the model estimates associations between variation in genome-wide markers and variation in phenotypic values. This means that the associations can be used to predict breeding values for non-phenotyped individuals which have genomic information. Such genome-based predictions have revolutionized many breeding programmes, enabling an efficient and early selection of candidates individuals and leading to significant genetic and economic gain per unit of time (Crossa et al., 2017; Hayes, Bowman, et al., 2009; Pryce et al., 2011).

Genome-based prediction is of particular interest in forest trees, as it could decrease the length of breeding cycles, which are long for such species, and cut the cost of phenotyping complex traits, such as drought tolerance and disease resistance (Grattapaglia & Resende, 2011, Isik 2014). Driven by the promising results obtained for simulations and first empirical approaches (Grattapaglia et al., 2011; Grattapaglia & Resende, 2011; Iwata et al., 2011), increasing numbers of experimental GS studies have been performed in recent years on many forest tree species (see Lebedev et al., 2020 for a recent review). Many of the studies have highlighted the attractiveness of genome-based prediction by reporting moderate to high prediction accuracies (Durán et al., 2017; Isik et al., 2016; J. Resende M. F. R. et al., 2012), and by reporting improved genetic gain per unit of time due to 20-50% shorter generation interval for GS (Chen et al., 2018; Lenz et al., 2017; Ratcliffe et al., 2015; Resende Jr et al., 2012).. However, this reduction in generation interval is also possible with pedigree-based prediction (if considering forward selection) and it is not clear if higher genetic gain per unit of time with GS is due to genome-based predictions enabling shorter generation interval, higher accuracy, or both. In fact, several studies in forest tree breeding report that pedigree-based predictions and genome-based predictions have similar accuracy to genome-based predictions (Beaulieu et al., 2014; Lenz, Nadeau, Azaiez, et al., 2020; Thistlethwaite et al., 2017, 2019; Zapata-Valenzuela et al., 2012, 2013; Zhou et al., 2020). Furthermore, use of an incomplete or error-containing pedigree tends to distort the comparison with genomic data (El-Dien et al., 2018; Li et al., 2019). Such errors may be common in forest tree breeding programmes and this penalizes pedigree-based evaluation (Doerksen & Herbinger, 2010; Munoz et al., 2014). The advantage of GS may therefore stem at least partly from the errors inherent to pedigree-based selection (Lenz, Nadeau, Azaiez, et al., 2020). Clarifying the conditions in which genome-based models can deliver real benefits remains a prerequisite for full exploitation of the advantages of GS in forest trees.

The access to within-family variability provided by molecular markers should increase the benefits of GS relative to pedigree-based selection, making it possible to improve the management of diversity. Indeed, accurate within-family prediction would facilitate the exploitation of within-family genetic variability rather than inter-familial variability, preventing the over-representation of certain lineages during selection and subsequent drift (Allier et al., 2019; Jannink, 2010; Rauf et al., 2010). However, little attention has been paid to the accuracy of genome-based prediction within families in forest trees, mostly due to the limited number of individuals per half- and full-sib family generally used in progeny trials. This issue has been addressed in only a few studies. Fuentes et al. (2017) and Cros et al. (2019) each studied a single large full-sibling family, making it difficult to extrapolate their results to more general cases. In three other studies (Pégard et al., 2020; R. T. Resende et al., 2017; Ukrainetz & Mansfield, 2019), the accuracies of within-family predictions were substantial, but variable.

Maritime pine (*Pinus pinaster* Ait.) covers 4.2 million hectares in south-western Europe (Abad Viñas et al., 2016). A breeding programme based on a recurrent selection scheme was initiated for this species in France in the 1960s (C.-E. Durel, 1992, GIS 2002). Starting from a base population (600 G0 individuals) selected for growth, environmental adaptation and stem straightness, two breeding cycles were performed using estimated breeding values from pedigree-based model (Bouffier et al., 2016). The potential of genome-based prediction in maritime pine breeding has already been highlighted in two previous studies (Bartholomé et al., 2016; Isik et al., 2016), but, as for most forest tree species, it is essential to investigate in greater depth the conditions in which genome-based prediction is clearly superior to pedigree-based one.

The aim of this study was to evaluate the ability of genome-based prediction to capture Mendelian sampling term in a maritime pine breeding population with empirical and simulation approaches. In both cases, the accuracy of genome-based prediction was estimated at the population as well as at the within-family levels and was compared with that of pedigree-based prediction. The real data were obtained for a population of 39 full-sib families with family sizes ranging from 10 to 40 individuals per family. The simulation, designed to mimic the conditions of the maritime pine programme, added other scenarios not observed in the real population, including variations of heritability, training set size and marker density.

## 2. Materials and Methods

### 2.1. Exploring accuracy of predictions with real data

#### 2.1.1. Maritime pine trial

A maritime pine trial was established in 2011 in the Landes de Gascogne forest at Le Barp (Lat 44.62, Long -0.77). A complete block design was used, with 89 full-sib families and 10 checklots, each containing 48 individuals planted in six-tree plots (1,250 trees/ha). The full-sib families considered were taken from the third generation of the French maritime pine breeding programme (i.e., the pedigree of the trees was known to grandparent level).

Preliminary simulations were performed to determine the most appropriate proportion of families and offspring per family from the total trial population to maximize GS accuracy at both global and within-family levels (**Appendix 1**). Based on the simulation, an optimal sample of 40 families was obtained, 30 of which contained 20 individuals each and 10 of which contained 40 individuals. The selected families were representative of the genetic diversity present in the trial and the within-family samples were representative of the phenotypic variability of each family. The largest families were chosen to represent a contrast in terms of relatedness with the rest of the sampled population. The larger families corresponded to five of the best-related and five of the worst-related families (referred to hereafter as “well-related” and “poorly related” families). The average relatedness to the rest of the population was 0.03 for the well-related families and 0.01 for the poorly related families (calculated from pedigree data). Considering families with large numbers of offspring is key to investigating within-family predictive ability. After genotyping, our study set, called POP_R,_ contained 833 individuals (see Results) with an effective population size of 25 (Lindgren et al., 1996). Thirty-nine families could be used to assess within-family predictive ability, including nine families of more than 30 individuals each.

#### 2.1.2. Genomic and pedigree information for POP_R_

Genomic DNA was extracted from young needles collected from each individual of POP_R_. DNA quantification and quality control were performed by fluorimetry (Qubit 2.0, Life Technologies, Thermo Fisher Scientific, USA) and spectrophotometry (NanoDrop Technologies, Wilmington, DE, USA), respectively. Genotyping was performed by Thermo Fisher Scientific (Thermo Fisher Scientific, Santa Clara, CA, USA) with the 4TREE Axiom 50K single-nucleotide polymorphisms (SNP) multi-species array (Guilbaud et al., 2020). Samples with a call rate of less than 97% were excluded from further analysis. In addition to the quality control filters at SNP level (CallRate ≥ 85%, fld-cutoff ≥ 3.2, het-so-cutoff: ≥ -0.3) suggested by Thermo Fisher Scientific, we also excluded SNPs with more than 5% Mendelian segregation errors, SNPs with a repeatability below 98% (estimated with 42 duplicated samples), and SNPs with a minor allele frequency (MAF) below 1%. Finally, 833 individuals characterized for 8,235 SNPs were available for this study. Missing genotypes were imputed by assigning the average genotype within the full-sib family. It was not possible to apply more sophisticated imputation methods due to the lack of a genetic map. We computed a realized genomic relationship matrix (**G**) following Van Raden (2008), with the R package AGHmatrix (Amadeu et al., 2016):

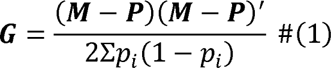

where ***M*** and ***P*** are matrices of dimensions *n* (number of individuals) x *p* (number of markers). ***M*** gives the genotype at each locus, coded -1 for one of the homozygotes, 0 for heterozygotes and 1 for the other homozygote, and the ***P*** is a matrix of allele frequencies expressed as 2(*p_i_* - 0.5), where *p_i_* is the observed allele frequency at marker *i* for all genotyped individuals.

Pedigree information was also available for POP_R_ at the parental (41 seed parents and 40 pollen parents) and grandparental (103 initial progenitors from the base population of the breeding programme) levels. Pedigree errors were detected with the R package pedtools (Dehli Vigeland, 2022), by comparing the genotyping data of POP_R_ with the genotyping data available for 78 of 81 parents. Pedigree errors were detected for 42 individuals (5% of individuals), i.e. when more than 1% of mismatches were detected, for all SNPs, between these individuals and their theoretical parents. Non-genotyped parents were assumed to be correct. Where possible, pedigree errors were corrected by identifying a new parent with less than 1% mismatches with the descendant. Thus, 27 individuals were reassigned to another sampled family, decreasing the total number of families from 40 to 39; the other 15 individuals were considered to be of unknown parentage. Within-family predictive ability was assessed for 39 full-sib families; nine large families with a mean of 34 individuals per family (30 to 40) and 30 families with a mean of 17 individuals per family (10 to 20). A complete corrected version of the pedigree was used to calculate an additive relationship matrix **A**, for a total of 1014 individuals.

#### 2.1.3. Phenotypic data

All individuals in the original trial, including those of POP_R_, were phenotyped at the age of eight years for height (HT) and deviation of the stem from verticality (DEV). Phenotypic values were adjusted for within-site spatial effects with spline functions implemented in the R package breedR (Muñoz & Sanchez, 2020) and pedigree information available for the whole trial. Below, we consider only these adjusted phenotypes for POP_R_. Heritability for HT and DEV within POP_R_ were 0.13 and 0.21, respectively, for estimates based on genomic data, and 0.17 and 0.25, respectively, for estimates based on pedigree information.

#### 2.1.4. Genome-based and pedigree-based models

Breeding values were estimated for each trait and for the n individuals using the model:

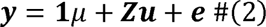

where ***y*** is the vector of adjusted phenotypes (dimension *n* x 1), **1** is the vector of 1s, *µ* is the population mean associated with a vector **1** of dimension (*n* x 1), ***Z*** is the incidence matrix (dimension *n* x *n*) connecting the phenotypes to the vector of breeding values ***u*** (dimension *n* x 1) and ***e*** is the vector of residuals (dimension *n* x 1). The ***u*** and ***e*** are assumed to be independent from each other and to follow a normal distributions of the form 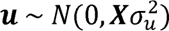 and 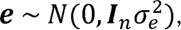 where ***X*** is either the genome-based (realized) relationship matrix ***G*** or pedigree-based (expected) relationship matrix ***A***, 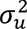 is the associated variance of breeding values, ***1***_*n*_ is the n-dimensional identity matrix and 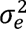 is the variance of residual effects. Mixed-model equations were solved to predict the random genetic effects ***u***:

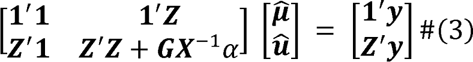

where ***X***^-1^ is the inverse of ***X*** and 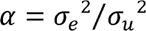 (Henderson, 1975; Mrode & Pocrnic, 2023). These two model versions (GBLUP and ABLUP) respectively gave us genome-based (GEBV) and pedigree-based (EBV) estimates / predictions of breeding values. All model fitting was performed with the R.4.2.2 environment (R Core Team, 2022) using the R package breedR (Muñoz & Sanchez, 2020).

#### 2.1.5. Cross-validation scenarios and assessment of prediction accuracy

Two cross-validation scenarios (CV) were used to assess the prediction accuracies of the GBLUP and ABLUP models (**Fig.1**): CV1 was used to assess within-family accuracy for the 39 families of POP_R_, whereas CV2 focused on the nine large families (i.e., the families from which 40 individuals were sampled). In the **CV1** scenario, the training set (T_set_) included 40% of each family and the validation set (V_set_) included the remaining 60% of each family. The **CV2** scenario was divided into six subscenarios. For the first subscenario, the T_set_ consisted of POP_R_ minus the large families and the V_set_ included all the individuals from large families. The number of individuals from large families included in the T_set_ was progressively increased in the other five subscenarios (5, 10, 15, 20 and then 25 individuals per large family), with a corresponding reduction of the contribution of other families to T_set_ so as to maintain a constant T_set_ size (the individuals from non-large families not included in the T_set_ became part of the V_set_, thereby also keeping the size of this set constant).

**Fig.1.**
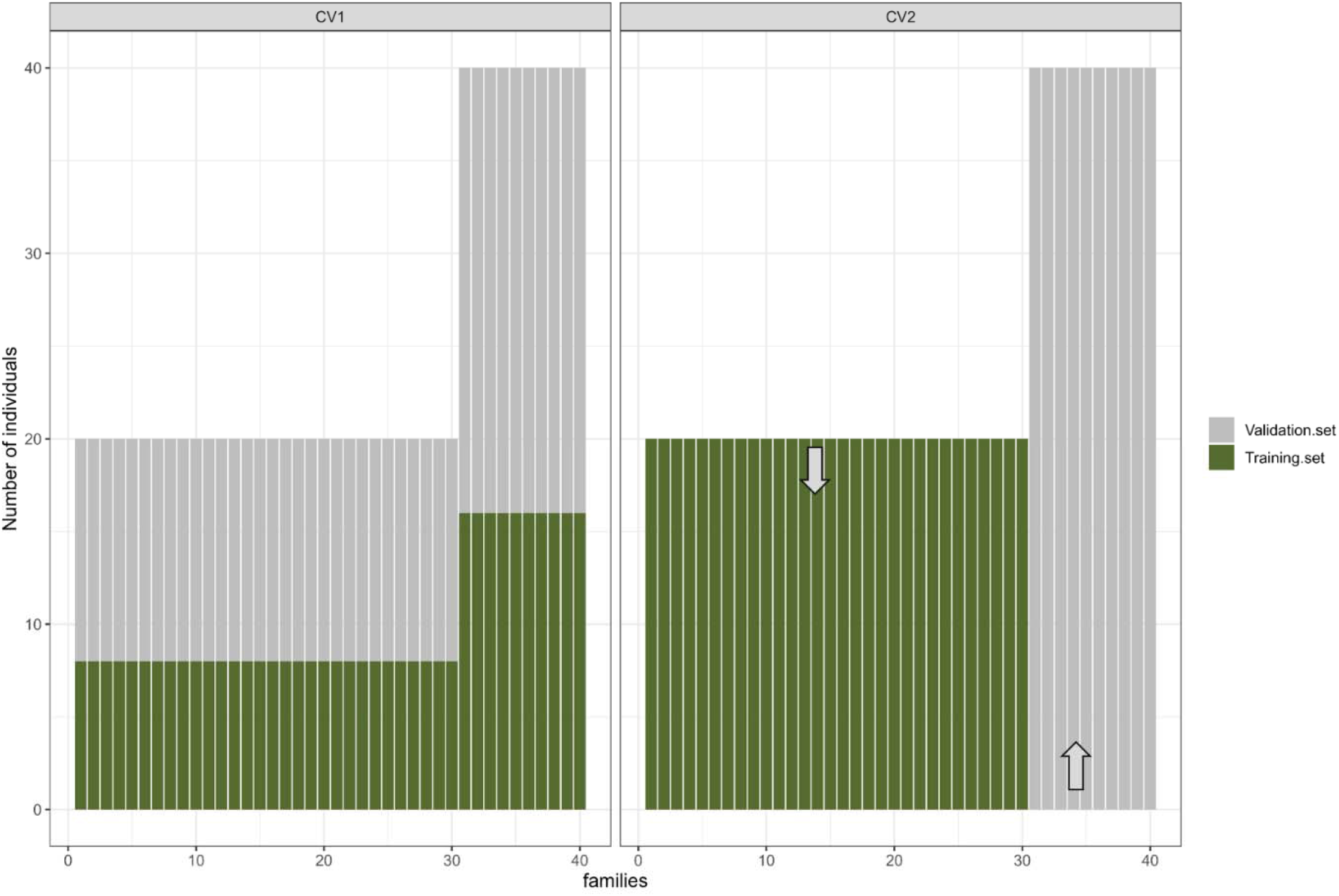
Cross-validation scenarios CV1 and CV2 performed with ABLUP and GBLUP models. In CV1, the training set include 40% of each family, the remaining 60% being the validation set. In the first sub-scenario of CV2, all non-large families constitute the training set and all large families constitute the validation set. For other sub-scenarios of CV2, the contribution of non-large families to the training set decreases in favor of large families. Training sand validation set sizes remain constant between sub-scenarios of CV2.

Each cross-validation subscenario was replicated 100 times. For each replicate, predictive ability was calculated by determining Pearson’s correlation coefficients for the correlation between adjusted phenotypes (*y*) and (G)EBV (*ŷ*) for individuals in the V_set_. Predictive ability was calculated at global level (for all individuals in the V_set_) or within families (considering each family separately). Note that within-family prediction is only meaningful for the GBLUP model. Namely, ABLUP model will only predict parent average component of the breeding value for non-phenotyped full-sibs, meaning that the resulting EBV will be the same for all full-sibs giving a within-family predictive ability of 0.

We calculated prediction accuracy, defined as the correlation between true (*g*) and predicted breeding values (*ĝ*), at the global level, to facilitate comparison with other studies, which frequently report this metric (Legarra et al., 2008):

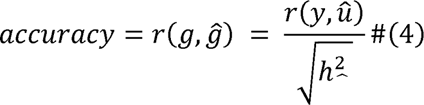

where h_^2_ is the heritability of the trait. We retained predictive ability as the metric for within-family analyses because dividing by the heritability defined at population level gives rise to values much higher than 1 or lower than -1. Instead, we focused on the deviation from 0.

### 2.2. Identifying key parameters for prediction accuracy with simulations

#### 2.2.1. Description of the simulation model

Stochastic simulations based on an allelic model were performed with the R package AlphaSimR (Gaynor et al., 2021). Briefly, from a base population representative of the maritime pine genome (Chagné et al., 2002; Chancerel et al., 2013; Jaramillo-Correa et al., 2020; Milesi et al., 2023), we simulated successive breeding populations based on a single trait (equivalent to HT), taking into account the characteristics of the real French maritime pine breeding program. Details of the simulations are provided in **Appendix 2** and the code is available at https://github.com/HighlanderLab/vpapin_pine_gs. Simulated phenotypes and genotypes for the final population, POP_S_ (the simulated version of POP_R_), were used to fit GBLUP and ABLUP models as described for the real data. The analyses were performed for the CV1 scenario (40% of individuals in each family included in the T_set_). Heritability of simulated phenotypes was set to 0.13 and the number of markers used was 8235, to mimic the real-life data as closely as possible. Ten independent replications of the entire process described above were performed to ensure that the results were robust.

#### 2.2.2. ABLUP and GBLUP prediction accuracy under different scenarios

Stochastic simulations were used to extend the comparison of prediction accuracy between GBLUP and ABLUP to different scenarios in which trait heritability (h²), training set size (nT_set_) and the number of markers (nSNP) were varied. For this purpose, POP_S_ was extended by generating 100 individuals for each of the 40 initially sampled families. Accuracy was assessed with a unique cross-validation scenario similar to CV1 (all families contributing equally to the T_set_) but with a fixed-size V_set_ of 1,200 individuals (evenly distributed between families). The values taken by the three parameters mimic real possibilities for maritime pine breeding:

- The size of the T_set_ was set to nT_set_ = 400, 600, 1600 or 2600 individuals, corresponding to 10, 15, 40 and 65 individuals per family, respectively. Such numbers are usually available in most forest tree breeding programmes. These numbers are also economically viable, because they require only 10, 15, 40 and 65% of the population to be phenotyped.
- The number of markers was set to nSNP = 8235, 17220 or 35000 SNPs, corresponding to marker densities of 5.7, 12 and 24 markers/cM, respectively. Data for 8235 SNPs are already available in our real maritime pine dataset, but this number could be increased by developing chips with a higher density of markers.
- The heritability of the trait was set to *h^2^* = 0.13, 0.33, or 0.50. Mean heritability for HT in the maritime pine breeding programme is 0.33, but HT ranges between 0.13 and 0.50, depending on the trial considered.

## 3. Results

### 3.1. Global and within-family predictive ability for maritime pine data

#### 3.1.1. Global prediction accuracies of GBLUP and ABLUP

Global prediction accuracies were estimated with the CV1 scenario (**Fig.2**). Mean prediction accuracy was similar for the two traits at 0.52 (±0.08) and 0.55 (±0.07) for HT and DEV, respectively. Mean prediction accuracy was slightly higher for the GBLUP model than for the ABLUP model, at +0.05 (±0.11) for HT and +0.02 (±0.10) for DEV, but the differences were significant only for HT. For this CV1 scenario, we also varied the percentage of individuals from each family included in the T_set_. The proportion included ranged from 20% to 80%, and accuracy increased with this percentage, from 0.45 (±0.09) to 0.62 (±0.18), but in a similar manner for ABLUP and GBLUP. We present only the data for a proportion of 40% (the mode) in **Fig.2**.

**Fig.2.**
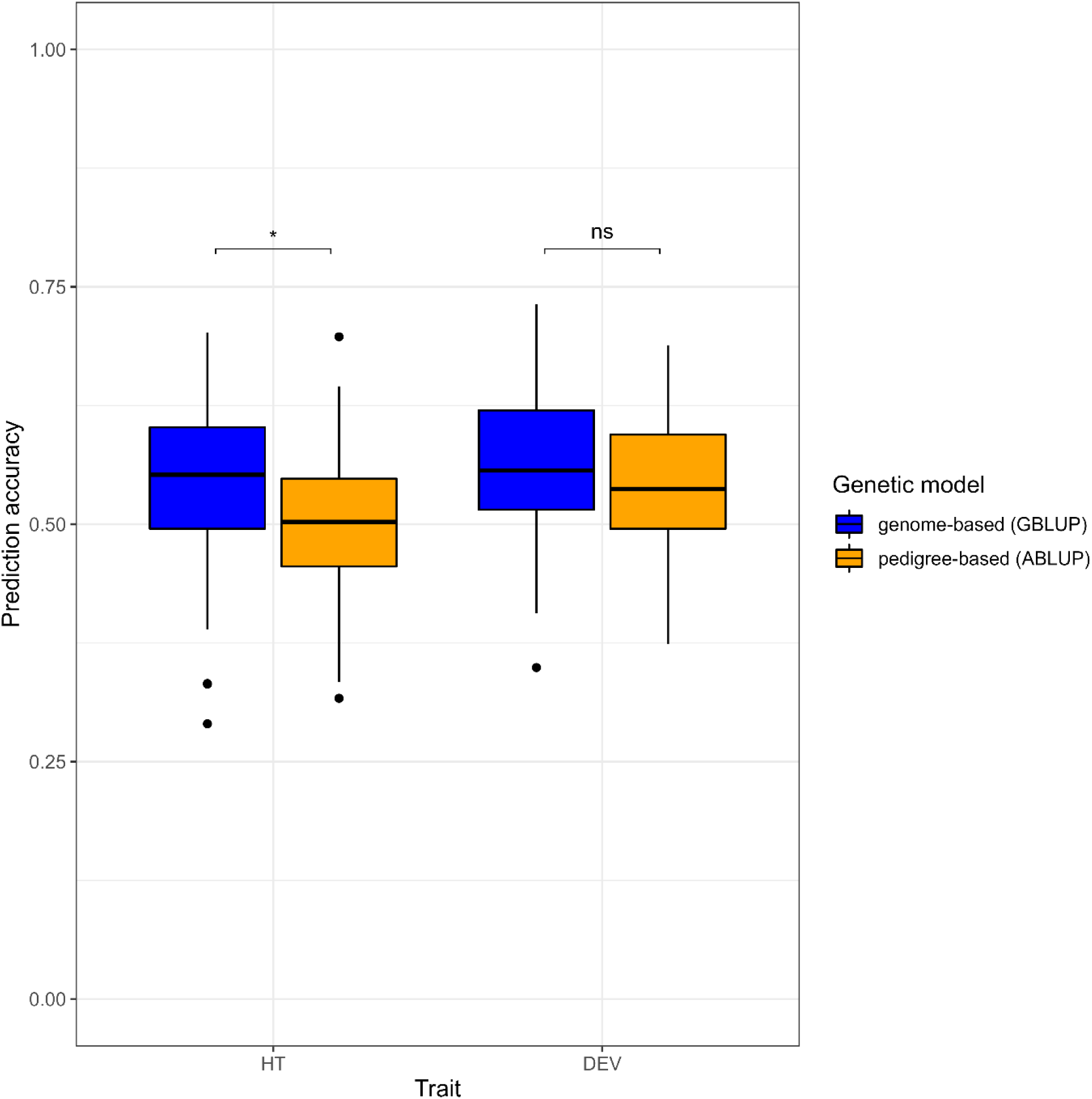
Global prediction accuracies obtained in the scenario CV1 with ABLUP and GBLUP models, for height (HT) and stem deviation to verticality (DEV)

In the CV2 scenario, global accuracy varies greatly with the structure of the T_set_ (**Fig.3**). Adding more individuals from the large families and removing individuals from the other families increased mean accuracy, with a maximum value of 0.68 (±0.10) obtained for both traits when 10 to 15 individuals from large families were included in the T_set_. Global accuracy declined if the number of individuals from large families was increased any further. In other words, accuracy was highest when all families were equally represented in the training set, and the overrepresentation of large families did not improve prediction. In most subscenarios, the differences between the GBLUP and ABLUP models were not considered significant, given the similarity of the mean values and the large overlap of their standard deviations.

**Fig.3.**
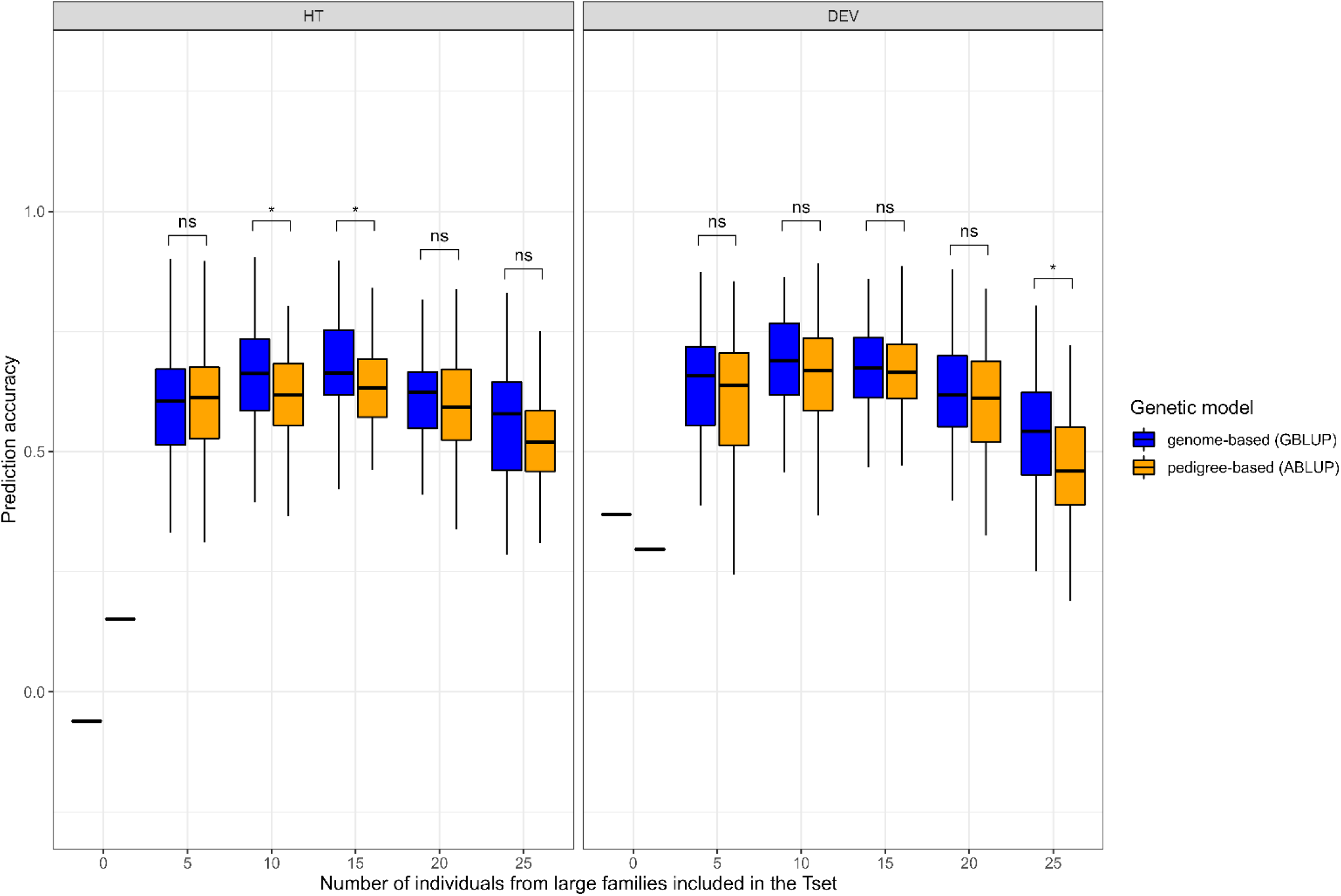
Global prediction accuracies obtained in the different sub-scenarios CV2 with ABLUP and GBLUP models, for height (HT) and stem deviation to verticality (DEV)

#### 3.1.2. Within-family genomic prediction accuracy

The large number of individuals per family in our design made it possible to assess within-family predictive ability. **Fig.4** shows the predictive abilities obtained with the CV1 scenario when 40% of the individuals from each family were included in T_set_, for each of the two traits (**Fig.4** A and B). Within-family predictive ability was therefore estimated for the 60% of individuals per family included in the V_set_ (a mean of 20 individuals for each of the nine large families and 10 individuals for the other 30 families). For each family, the variance of prediction accuracies was very high, indicating that the choice of individuals in the T_set_ (and therefore in the V_set_) had a major impact on prediction accuracy. Mean prediction accuracy differed between families (**Fig.4**), ranging from -0.43 (±0.20) to +0.45 (±0.18) for HT and from -0.35 (±0.24) to +0.46 (±0.22) for DEV. These mean values were centered on 0, with a very low cross-family mean (-0.01 (±0.33) for HT and +0.06 (±0.33) for DEV). Despite the similar distribution of within-family predictive ability for the two traits, the ranking of families differed significantly between HT and DEV (Kendall correlation of +0.01).

**Fig.4.**
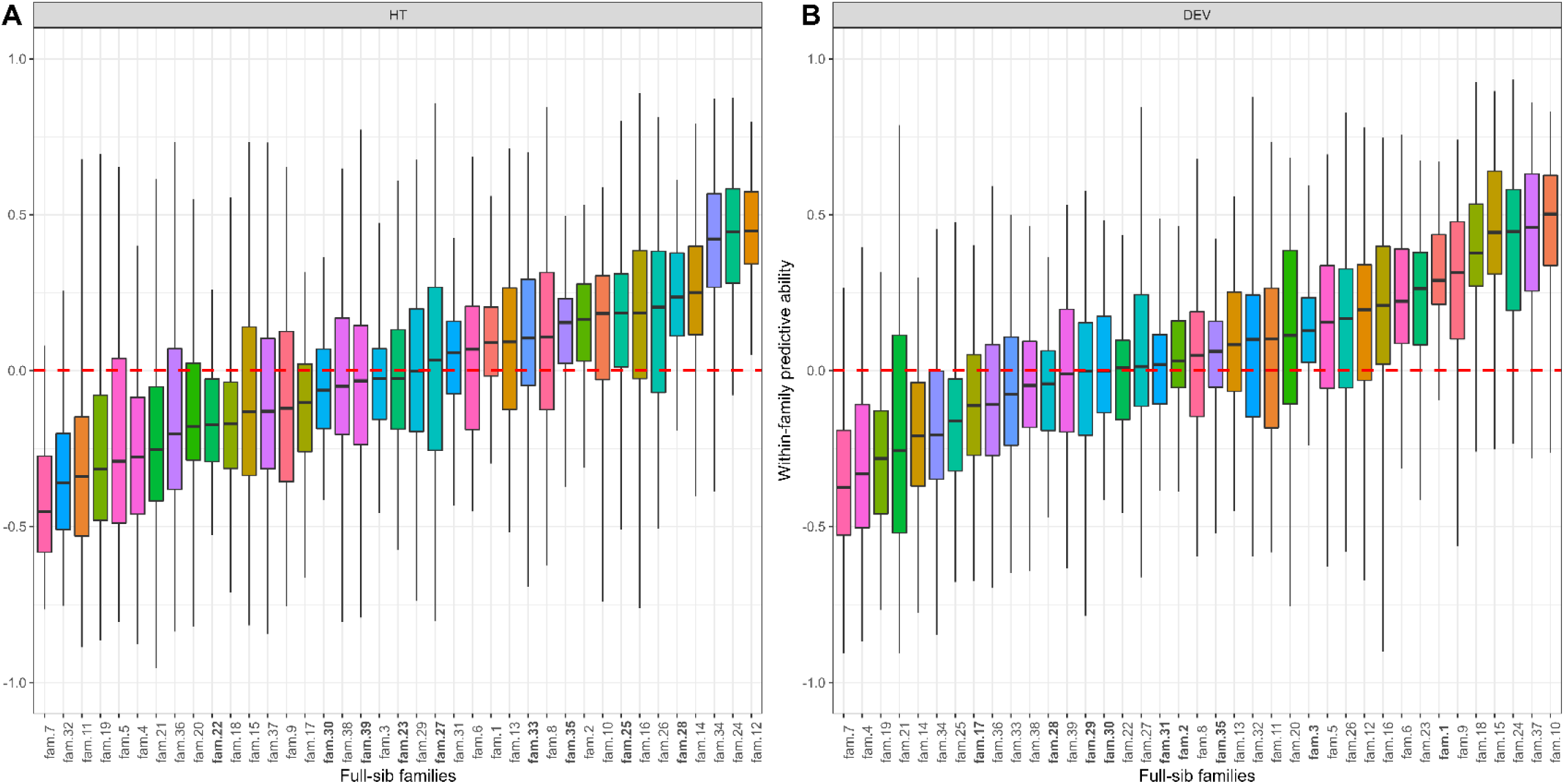
Genome-based within-family predictive ability obtained in the scenario CV1 for each of the 39 full-sib families, for height (HT) and stem deviation to verticality (DEV)

We then investigated within-family predictive ability further, using the CV2 scenario (**Fig.5**). Once again, the variance of prediction accuracy increased significantly with decreasing V_set_ size. Furthermore, the ranking of families on the basis of prediction accuracy differed between the two traits. Prediction accuracy was low for both traits, but prediction accuracy was higher for families well-related to the training set than for poorly related families (mean +0.12 (±0.08) for HT and +0.16 (±0.09) for DEV). For well-related families, adding more individuals to the T_set_ either had no impact on within-family mean prediction accuracy, or it increased this accuracy, as observed for HT in families 1, 2 and 28. For families poorly related to the training set, adding more individuals to the T_set_ had no impact, increased or, more surprisingly, decreased within-family predictive ability, as observed for both traits in family 22. For both scenarios (CV1 and CV2), within-family predictive abilities remained close to 0 on average, despite strong variation between families, but could potentially explain the equivalence in terms of global prediction accuracy between the GBLUP and ABLUP models.

**Fig.5.**
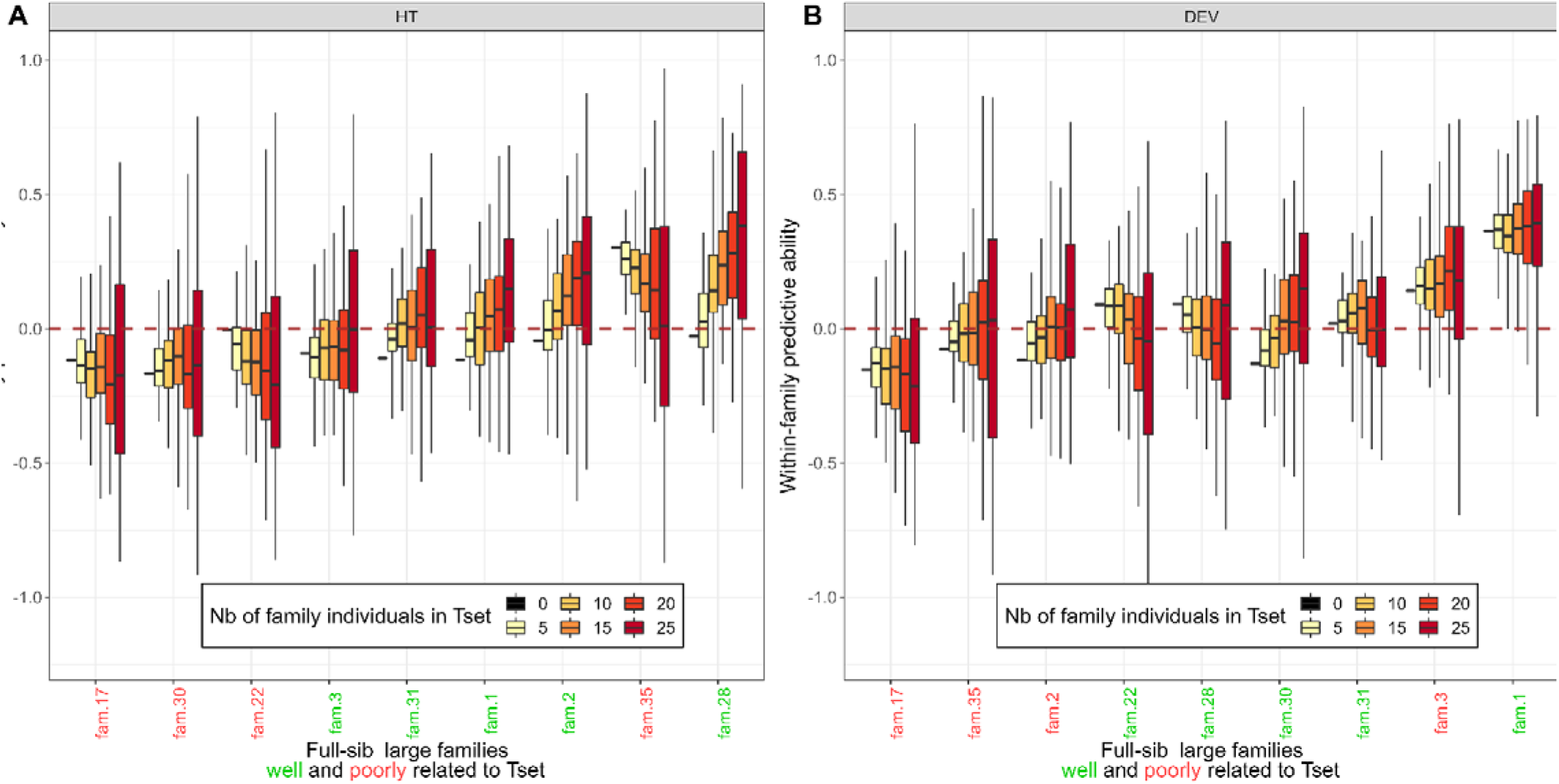
Genome-based within-family ability obtained in the sub-scenarios CV2 for each of the 9 large full-sib families, for height (HT) and stem deviation to verticality (DEV)

### 3.2. Use of simulations to explore prediction accuracy

#### 3.2.1. A relevant simulation model

The comparison of simulated and real data is an interesting preliminary step in assessments of the relevance of the simulation model been constructed. For both models, GBLUP and ABLUP, the simulated and real data gave very similar prediction accuracies (**Fig.6**) in terms of the mean (0.54 (±0.07) and 0.55 (±0.08) for real and simulated data, respectively, with the GBLUP model, 0.49 (±0.07) and 0.47 (±0.07), respectively with the ABLUP model), and the distribution of values (the coefficient of variation for prediction accuracy being 15% and 14% for real and simulated data, respectively, with the GBLUP model, and 15% and 16%, respectively, with the ABLUP model). With the simulated data, prediction accuracy was slightly better for GBLUP than for ABLUP. The agreement between the simulated and real data was very high when accuracy comparisons were made within families (**Fig.7**). The high variance of the within-family predictive abilities in each family reported previously on the basis of real data was also observed with the simulated data, as were the large differences between families in terms of mean within-family predictive ability. Within-family predictive ability ranged from -0.43 (±0.20) to +0.45 (±0.18) with the real data, and from -0.31 (±0.27) to +0.42 (±0.28) with the simulated data. The ranking of families on the basis of within-family accuracy differed between the real and simulated data, as the genotypes were simulated independently of the real data.

**Fig.6.**
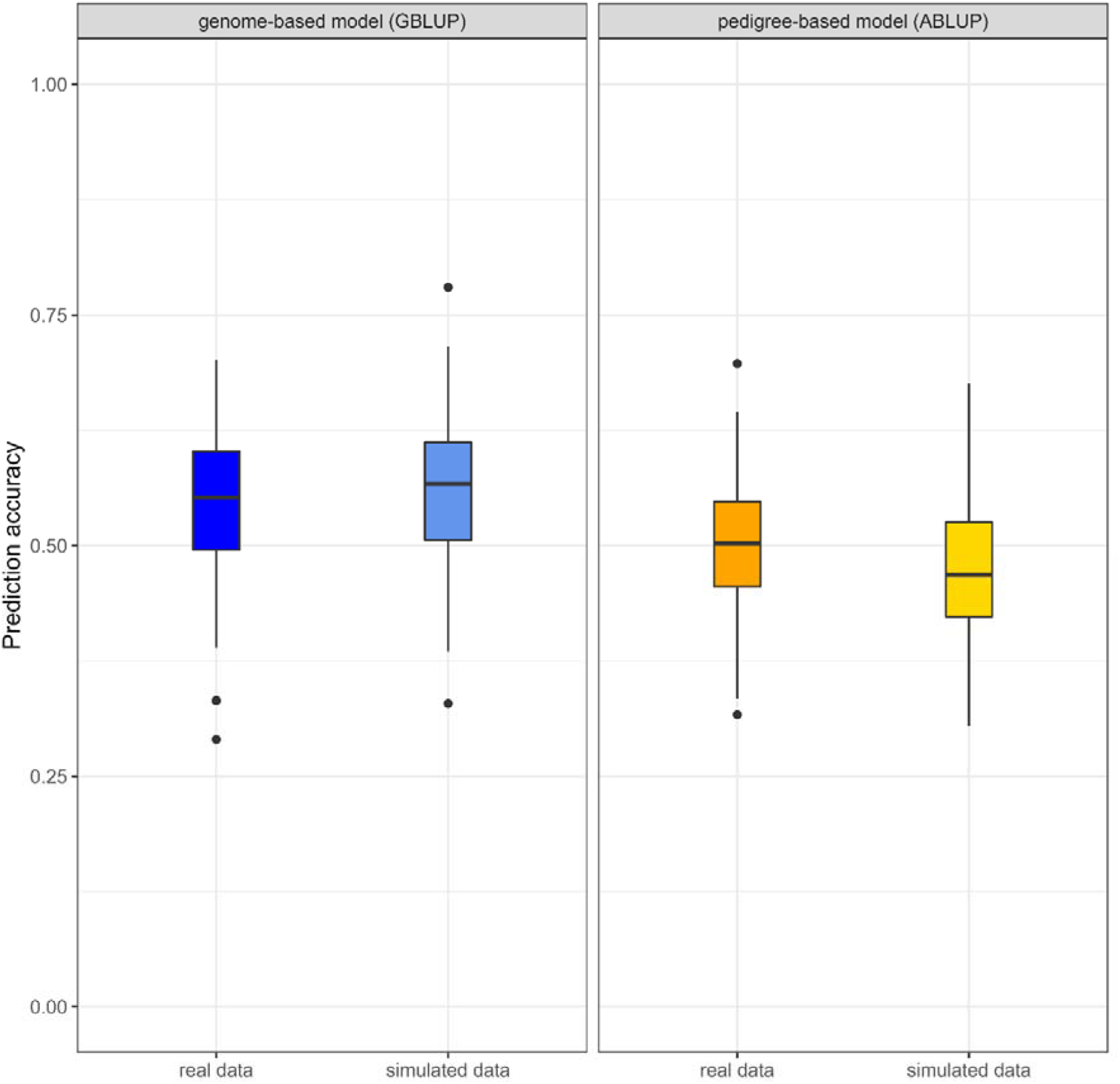
Global prediction accuracies obtained in the scenario CV1 for height with ABLUP and GBLUP models based on real or simulated data.

**Fig.7.**
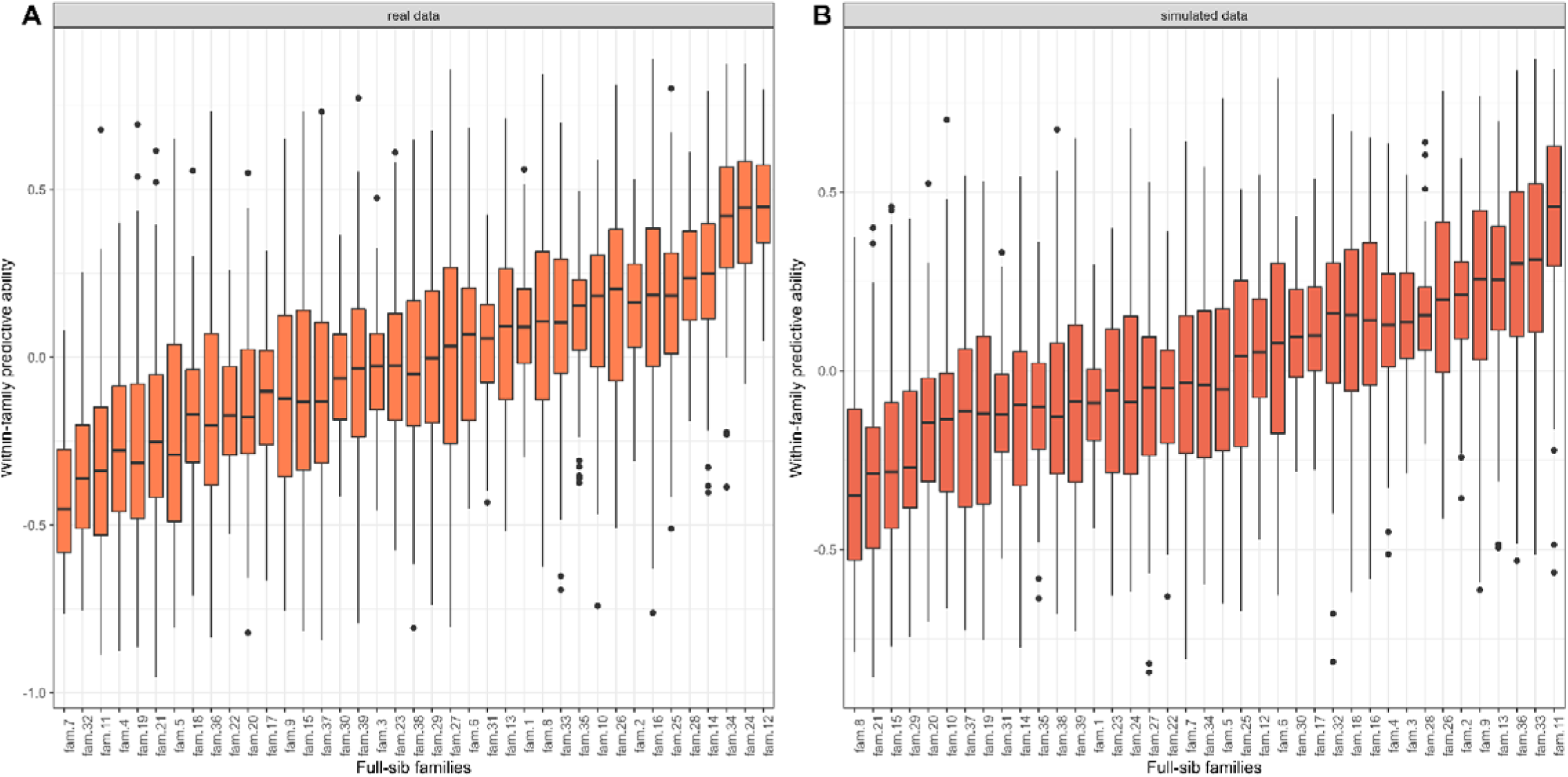
Genome-based within-family predictive ability obtained in the scenario CV1 for height (HT) for each of the 42 full-sib families with real or simulated data.

#### 3.2.2. Determining relevant conditions for GS implementation

Starting from the initial conditions defined by the real data (h²=0.13, nSNP=8235 and nT_set_ *ϵ* [167:667] depending on the CV1 subscenario considered), new scenarios with different conditions were explored through simulations. **Fig.8** shows the overall prediction accuracy for different combinations of h², nT_set_ and nSNP. The objective was not only to study the behavior of accuracy as a function of the variation of these key parameters, but also to compare it with the accuracy of predictions based solely on pedigree, the reference in many forest tree improvement programmes.

**Fig.8.**
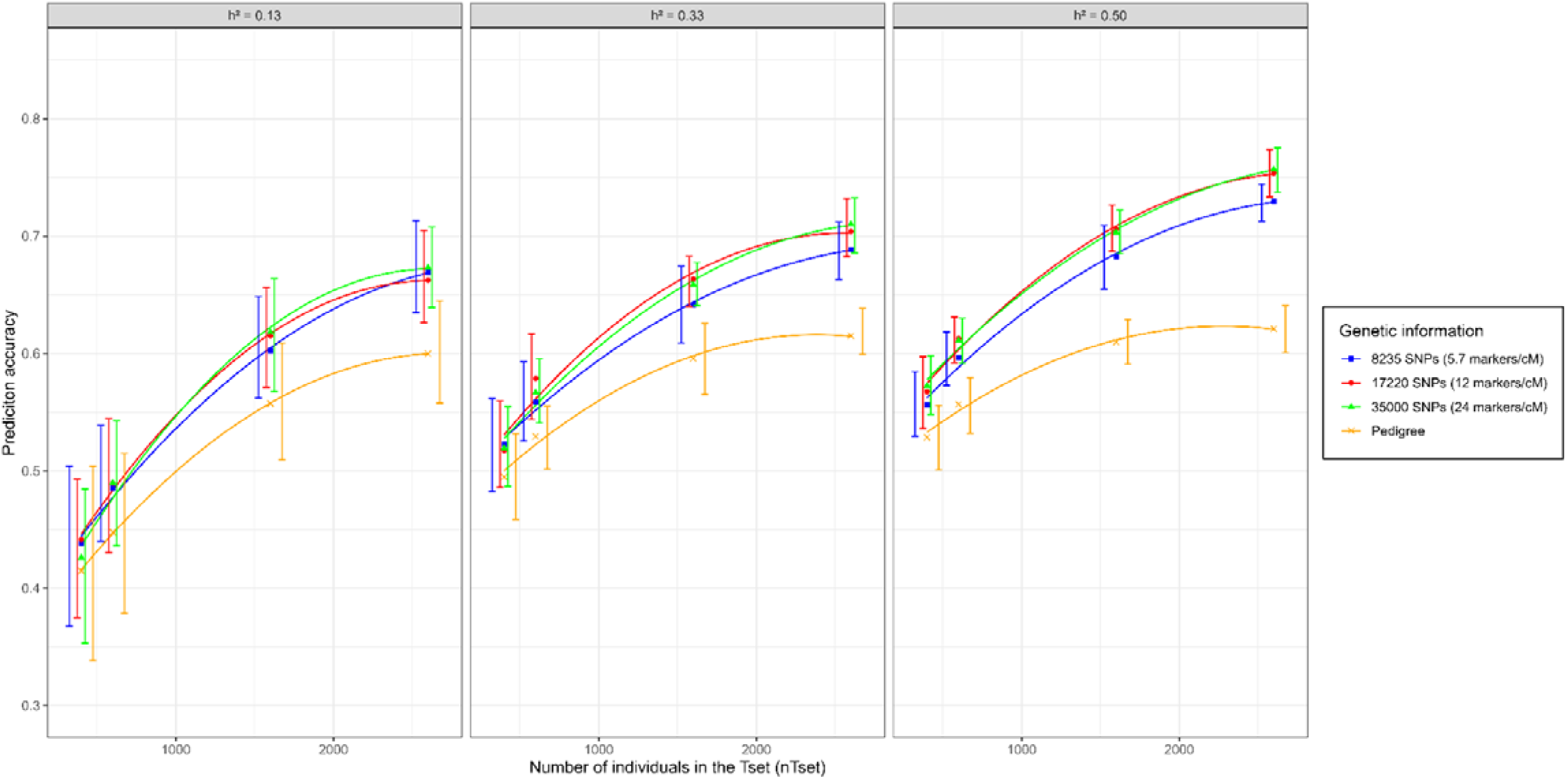
Global prediction accuracies for height (HT) with simulated data for different combinations of heritabilities (h²), training set sizes (nT_set_) and marker densities (nSNP)

Marker density did not appear to be of a critical importance for GS accuracy, as curves associated to genomic models (blue, red and green) overlapped in most situations. The greatest benefits of a higher marker density (12 and 24 markers/cM) in the tested scenarios were observed in situations in which a large T_set_ was combined with high heritability. A similar combination of parameters also resulted in a lower level of variation in accuracy. Similar results were obtained for the highest two densities, suggesting that the response is saturated when there are more than 17,000 markers.

Regardless of heritability, prediction accuracy appeared to be highly dependent on T_set_ size. GS prediction accuracy increased steadily for the first few increases in T_set_ size (nT_set_=400, 600, 1600) and then tended to stabilize at about nT_set_=2600, reaching a plateau with inflexion points between nT_set_=1600 and nT_set_=2000. The prediction accuracy of ABLUP models followed a similar pattern, but the advantage of GBLUP models over ABLUP models was greatest for larger T_set_ sizes and higher heritability. For a trait of average heritability (h²=0.33), the advantage of GBLUP models in terms of mean prediction accuracy was only +0.02 (±0.08) when nT_set_=400, whereas it reached +0.08 (±0.05) when nT_set_=1600 (+0.13 (±0.03) with 2600). The simulation results clearly indicate that the conditions under which the real data were analyzed were not optimal for revealing any advantage of the GBLUP model. According to the simulation data, the GBLUP model would be much more advantageous at higher nT_set_ (nT_set_≥1600) values, for traits of intermediate or higher heritability and with a larger number of markers.

For two of the situations in which GBLUP performed significantly better than ABLUP — with a minimal difference in one case (+0.06 (±0.05) in mean accuracy) and for the maximum observed difference in the other (+0.13 (±0.03)), we present within-family predictive abilities in **Fig.9**. As for all the within-family predictive abilities presented above, the variance was high and mean values differed considerably between families. However, in this case, mean prediction ability values were positive for most families, ranging from -0.03 (±0.18) to 0.34 (±0.13) for the first situation (h²=0.33, nT_set_=1600, nSNP=17220) and from 0.07 (±0.14) to 0.50 (±0.12) for the second situation (h²=0.5, nT_set_=2600, nSNP=35000). Taking all the families into account, mean within-family predictive ability was 0.18 (±0.14) and 0.29 (±0.17), respectively, for the two situations described, indicating a clear advantage of genome-based approach over the reference value of 0 associated with the pedigree-based model. The advantage of the GBLUP model over the ABLUP model in terms of overall prediction accuracy coincides with non-zero within-family predictive ability. This advantage of the GBLUP models was increasingly evident for higher values of within-family predictive ability.

**Fig.9.**
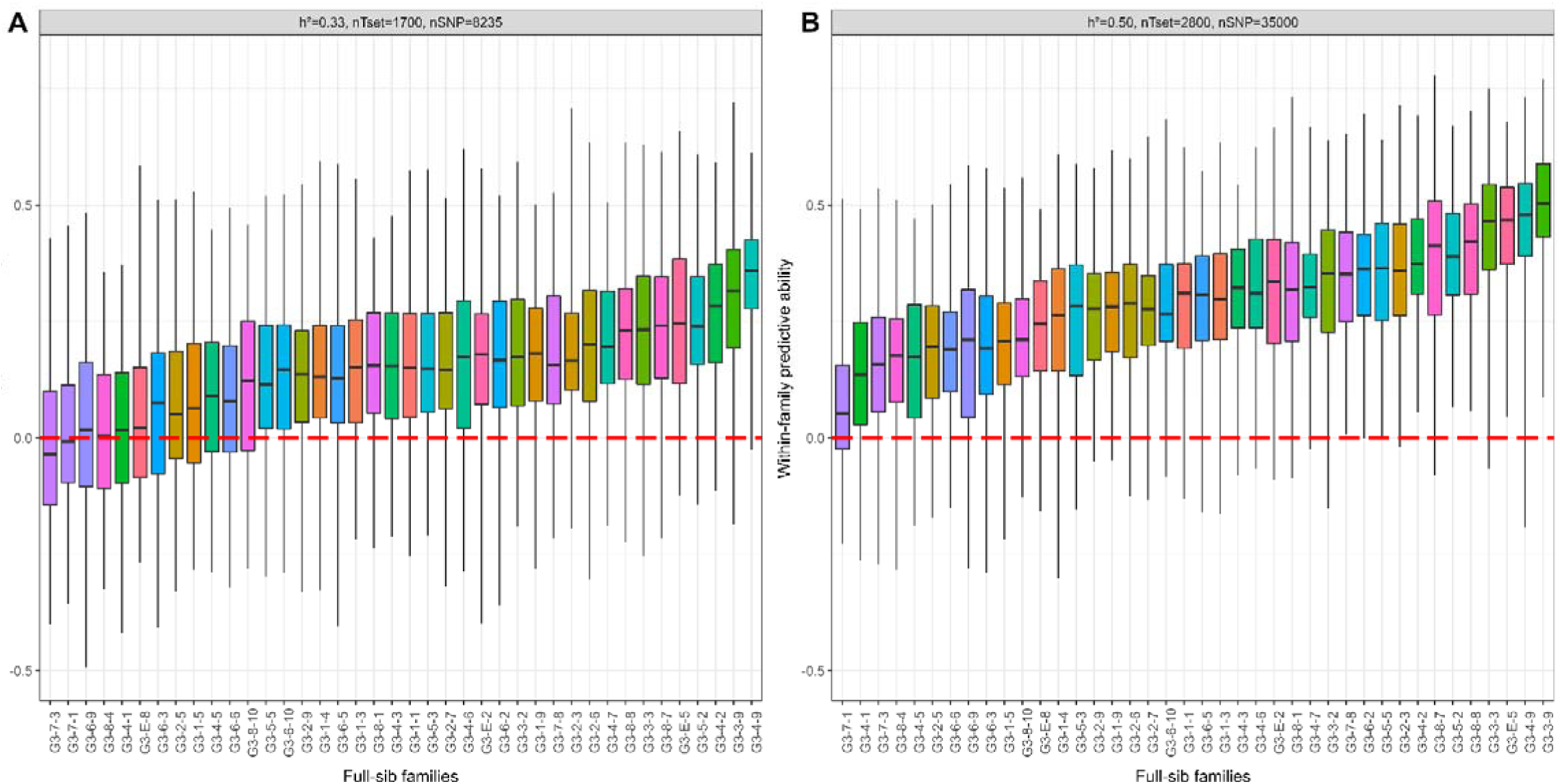
Genome-based within-family predictive ability for each full-sib family for height (HT), with simulated data and for two different combinations of heritability (h²), training set size (nT_set_) and marker density (nSNP)

## 4. Discussion

Before implementing genome-based prediction and selection in maritime pine and forest trees more generally, we need to clearly define the conditions in which the use of this approach has real advantages over conventional methods, particularly in terms of prediction accuracy (Bartholomé et al., 2016; Beaulieu et al., 2014). Unlike most previous genomic studies on forest trees, we evaluated genome-based prediction accuracies by comparison with the corresponding pedigree-based prediction accuracies, and we investigated within-family accuracy. The reference considered here is key as it allows a fair comparison between predictions based on genome and pedigree information available at the same time point (seedling stage) without the need to calculate a selection response taking the length of breeding cycles into account. We found that even with a test design that *a priori* favoured within-family variability, the advantage of GBLUP over ABLUP models was not clearly significant, and that within-family predictive ability was zero, on average, across families. Stochastic simulations mimicking our selection scheme showed that our real case was close to the tipping point for recommended training set size, beyond which genome-based predictions would start to show its full potential relative to conventional pedigree-based predictions.

### 4.1. Equivalence of ABLUP and GBLUP prediction accuracies with a maritime pine dataset

#### 4.1.1. Baseline for comparison of the ABLUP and GBLUP models

Bartholomé et al. (2016) reported a higher prediction accuracy for ABLUP than for GBLUP models in maritime pine, with similar conclusions drawn in several other studies on forest trees (Thistlethwaite et al., 2017; Zapata-Valenzuela et al., 2012; Zhou et al., 2020). An insufficiently high marker density has frequently been identified as the cause of a lack of advantage of genome-based predictions. Marker density was higher in this study than in previous studies in maritime pine (5.7 SNPs/cM versus 2.4 and 1.7 SNPs/cM in the articles by Bartholomé et al., 2016 and Isik et al., 2016, respectively). Furthermore, the original design of POP_R_, characterized by a limited number of full-sib families but relatively large numbers of trees per family, was expected to favor the genome-based approach, better capturing Mendelian sampling terms that drive the within-family variation.

However, GBLUP models had only a slight advantage over ABLUP models in terms of prediction accuracy for some of the cross-validation subscenarios evaluated (**Fig.2 and Fig.3**), and even this advantage may be due, at least in part, to the way in which prediction accuracy is calculated. Prediction accuracy, defined as the ratio of predictive ability to the square root of heritability of the predicted trait, requires the estimation of variance components, which are subject to additional errors (Legarra et al., 2008). As in other studies, GBLUP estimates of heritability were lower than the estimates obtained with the ABLUP model (El-Dien et al., 2018; Lenz, Nadeau, Mottet, et al., 2020; R. T. Resende et al., 2017). This resulted in higher prediction accuracies for the GBLUP model despite a predictive ability similar to that for the ABLUP model. However, predictive ability has been much less studied and poses problems of interpretation in terms of genetic gain because it includes environmental effects, justifying our choice to consider prediction accuracies.

We also decided to base our comparison solely on the accuracy, without converting it into a response to selection. The predictions obtained with pedigree-based model takes into account only the pedigree information for the individuals in V_set_ (not their phenotypic records). The predictions of this model are therefore available for use in early selection, as for genome-based predictions. We believe that the assumption of a shorter selection cycle for genome-based predictions does not hold and that this assumption introduces a bias, increasing the perceived advantage of GS over traditional approaches (Lenz et al., 2017; Ratcliffe et al., 2015; Resende Jr et al., 2012). Genome-based prediction is clearly advantageous when it enables pedigree recovery and correction (Zapata-Valenzuela et al., 2012), but similar results can routinely be obtained with a very small number of carefully chosen markers across the genome (Vidal et al., 2017). For the use of genome-based prediction to be considered truly advantageous, it must, therefore provide a higher accuracy than pedigree-based prediction associated with a complete corrected pedigree.

#### 4.1.2. Capture of linkage disequilibrium and relatedness in GBLUP models

Theoretically, the genome-wide markers used for genome-based predictions can provide additional information over and above that obtained from a simple pedigree. They can, for example, capture the effects of nearby QTL in linkage disequilibrium (Habier et al., 2007), and provide more accurate information about the relationship between any two given individuals (Nejati-Javaremi et al., 1997; VanRaden, 2008). However, these two advantages of marker use did not emerge clearly in this study. In our case, there are two principal explanations for the equivalence between ABLUP and GBLUP (**Fig.2 and Fig.3**), as outlined by other authors faced with similar results (Beaulieu et al., 2014). First, the estimation of genomic effects may not be very accurate, potentially due to the T_set_ being too small to guarantee a good estimate (Habier et al., 2013). For example, with simulation, Hayes et al. (2009) showed that the advantage of GBLUP over ABLUP models was apparent from 100 individuals per full-sib family. Second, it is possible that genome-wide markers capture only existing genetic relationships (Legarra et al., 2008), being much less effective at capturing actual linkage disequilibrium (LD) (Habier et al., 2007).

A lack of long-range LD has already been reported for conifers (Eckert et al., 2010; Kujala & Savolainen, 2012), including maritime pine (Isik et al., 2016; Plomion et al., 2014), suggesting that a large number of markers may be required for genome-based predictions (R. T. Resende et al., 2017; Thistlethwaite et al., 2019). Grattapaglia et al. (2011) performed simulations for forest trees that suggested that a density of about 10 markers/cM was required to increase the prediction accuracy of GBLUP to such an extent that it surpassed ABLUP when Ne=100. Most conifer genomes have a much smaller genetic map than physical size [1435 cM (Chancerel et al., 2013) and 24 Gb (Chagné et al., 2002), respectively, for maritime pine], suggesting that there is very little recombination in large parts of the genome. Thus, rather than the number of markers, it may be their distribution across the genome genome that counts. However, it can be difficult to improve the distribution of markers in the absence of a physical map for the species, as in maritime pine. The densities used in our real case would, therefore, be sufficiently high to capture pedigree-like relatedness, but they are probably still to low and the markers are probably not distributed appropriately to capture QTL information across LD. One alternative not considered here would be the use of alternative statistical approaches based on Bayesian variable selection, such as Bayes-B. Such methods have been shown to capture population LD more effectively and to give a greater weight to causal SNPs (Habier et al., 2007; Thistlethwaite et al., 2017), but their benefits are often case- and trait-specific, and may even disappear, particularly if the training population exceeds a certain size (Karaman et al., 2016).

#### 4.1.3. Genome-based within-family predictive ability

Unlike ABLUP model, GBLUP model capture realized relatedness between and within families through the **G** matrix. Nevertheless, regardless of the scenario tested here, within-family predictive ability for the real data was, on average, zero when all full-sib families were considered (**Fig.4**), indicating that underlying genome-wide marker associations in GBLUP (Stranden & Garrick, 2009) were not estimated accurately. For both traits, within-family predictive abilities were not significantly different from zero in most families, but there was considerable variation across the families, including some (15%) for which accuracy values were, surprisingly, significantly negative. For some relatively small families, the number of individuals included in the V_set_ was probably too small for a robust estimation of correlations, resulting in a very large standard deviation when all CV iterations were taken into account. By contrast, within-family predictive ability for large families was calculated with a mean of 20 individuals (in the V_set_), resulting in zero or slightly positive values, but with a lower standard deviation. Genome-based prediction accuracy was, therefore, mostly driven by capturing parent average term rather than capturing the Mendelian sampling term of breeding values. This partly explains the observed equivalence with pedigree-based prediction models, as suggested in other studies with similar results (Thistlethwaite et al., 2019).

Some specific scenarios of cross-validation imposing restrictions on relatedness between V_set_ and T_set_, particularly if V_set_ did not contain relatives of individuals present in T_set_, resulted in particularly low prediction accuracies at the global (**Fig.3**) and within-family levels (**Fig.5**). This suggests that relatedness was the main source of information in both GBLUP and ABLUP, with little or no additional information captured from LD to maintain prediction quality in the absence of relatedness. Globally, the relatedness between families in our population was low, with parents and grandparents at the founding level of the population producing a mean of 1.1 and 2.4 full-sib families, respectively. This structure may pose challenges that prevent GBLUP from outperforming ABLUP. In general, the use of a diversified training population is always desirable, to produce robust predictions and a validation set well related to the training set. The first condition could have been met by ensuring the equal representation of different grandparents and parents. The second condition is less easy to satisfy, as the siblings in V_set_ probably had few collaterals other than the remaining siblings already present in T_set_.

### 4.2. Use of simulations to identify conditions in which GBLUP has a better prediction accuracy than ABLUP

The stochastic simulation model yielded results similar to those obtained empirically, suggesting that we can have some confidence in the relevance of the results, in terms of both the trends observed and the absolute values obtained. The effect of increasing T_set_ size and the number of genome-wide markers on GS prediction accuracy was tested under three contrasting heritability scenarios, each with an explicit comparison with the prediction accuracy of ABLUP. These simulations showed that the size of T_set_ was the most important determinant of prediction accuracy in our study. The inflection point of the curves occurred somewhere between 1,500 and 2,000 individuals, consistent with the findings of deterministic approaches in other forest tree contexts (Grattapaglia et al., 2011; Grattapaglia & Resende, 2011). From this T_set_ size upwards, prediction accuracy begins to be significantly higher for GBLUP than for ABLUP. Increasing the number of individuals per family, as in the simulation scenarios in which T_set_ increased, made it possible to estimate breeding values with greater accuracy (Habier et al., 2013). Thus, the often-highlighted equivalence between GBLUP and ABLUP in terms of prediction accuracy for forest trees, regardless of the species considered, can be explained by the training sets, which rarely exceed 1000 individuals, being too small (Lebedev et al., 2020). By contrast, increasing the number of markers did not increase significantly the genome-based predictive accuracy, at least for traits with moderate heritability, suggesting that our initial marker density would have been sufficient if it had been coupled with a larger training population. This conclusion could be readily extrapolated to other conifers, which have similar genome sizes and effective breeding population sizes, but it would be more difficult to generalize it to deciduous species, which can have very different genome structures and LD profiles.

Complementary analysis of genome-based prediction accuracy based on deterministic approaches (Daetwyler et al., 2008) with empirical parameters for our population of maritime pine (see **Appendix 2**) were consistent with our stochastic simulation results.

Within-family prediction accuracy is rarely considered in genomic studies but appears to be key for the superiority of genome-based predictions over pedigree-based predictions to be expressed. Simulations showed that more accurate within-family prediction was associated with a greater accuracy advantage of GBLUP over ABLUP. For our study design, the suggested T_set_ size for efficient within-family prediction corresponds to between 40 and 65 individuals per full-sib family. This is a very important requirement for the implementation of genome-based prediction and selection, as the use of GBLUP models with zero within-family predictive abilities can have several negative consequences.

In the short term, the effectiveness of selection and the response to selection would be reduced if one of the sources of genetic variation in the population, the within-family variation due to Mendelian sampling, was not exploited. While this source of variation can be measured with genome-wide markers at the genotype level, attaching quantitative value to these genotypes requires sufficiently powered training set of phenotyped and genotyped individuals. Ensuring such sufficiently large training set is important to avoid longer-term consequence related to the fact that selection on underpowered genome-based predictions would be only leveraging variation between families. Such an approach would increase the risk of losing diversity due to the elimination of certain lineages and the co-selection of candidates from the same families. This loss of diversity due to a shift in the weighting between within-family and between-family selection would lead to long-term losses of genetic gain (Jannink, 2010) and an accumulation of inbreeding.

This tendency can be counteracted by selection methods based on the optimization of genetic contributions (Meuwissen, 1997; Woolliams et al., 2015) — so-called “optimal contribution selection” (OCS) — which allows a trade-off between short-term and longer-term gains through the application of constraints to the balance between parental genetic contributions (Gorjanc et al., 2018). Future studies should assess these optimal strategies which would, presumably, work better if genome-based predictions could discriminate between candidates within families more accurately (Hallander & Waldmann, 2009), with sufficiently large families in the training populations, thereby increasing the efficiency of selection and of the constraints imposed by the OCS.

## 5. Conclusion

Despite the undeniable potential benefits for forest trees, examples are lacking for which genome-based approaches have clearly demonstrated superiority over pedigree-based approaches. Using an ABLUP model with full and corrected pedigree information as a reference, we evaluated the accuracy of GS in a maritime pine trial with the largest number of individuals per full-sib family to date. Prediction accuracy was found to be similar for the pedigree-based and genome-based models, and within-family genomic accuracy for forward predictions was close to zero. By constructing a relevant simulation model, we were able to demonstrate that the number of individuals per family, and thus the overall size of the training set, is a key parameter for accurately estimating marker associations and for detecting a clear advantage of genome-based approach. This conclusion can be extended to many forestry contexts in which the equivalence between ABLUP and GBLUP prediction accuracies can be explained by suboptimal training set sizes and structures. Increases in training-set size may be readily achievable in forestry due to the large numbers of individuals commonly used in breeding programmes and decreasing genotyping costs. Effective within-family prediction, based on well-scaled genome-based approaches, will be key to maintaining diversity in the long term and ensuring genetic gain in the challenging years ahead.

## Abbreviations

ABLUP: pedigree-based best linear unbiased prediction
BLUP: best linear unbiased prediction
CV: cross-validation
DEV: stem deviation to verticality
DNA: deoxyribonucleic acid
EBV: pedigree-based estimated breeding value
GBLUP: genome-based best linear unbiased prediction
GEBV: genome-based estimated breeding values
GS: genomic selection
HT: height
LD: linkage disequilibrium
nT_set_: training set size
nSNP: number of SNP
OCS: optimum contribution selection
POP_r_: trees sampled for this study
POP_s_: simulated version of POP_r_
QTL: quantitative trait loci
SNP: single nucleotide polymorphism
T_set_: training set
V_set_: validation set

## Appendix

## Appendix 1: definition of sampling within the trial for genome-based analysis

The initial maximum capacity for genotyping in this study was 800 individuals. Thus, preliminary simulations were performed to determine a relevant choice of these 800 individuals among the total population in the trial.

### Choice of sampling scenario and families

We compared 3 sampling scenarios to select 800 individuals among the 89 families present in the trial: 40 families with 20 individuals each (A), 32 families with 25 individuals each (B) and 20 families with 40 individuals each (C). The best scenario was selected on the basis of the prediction accuracy, following a three-step procedure that was replicated 10 times independently: 10 times independently:

- Genotypes and phenotypes were simulated for the families present in the trial using the MoBPS software (Pook, 2022).
- For each scenario, the subset of 20, 32 or 40 families was selected with the goal to minimize the average genomic relationships among the subset (with genomic relationships calculated using the simulated genotypes) and with a simple trail-and- error optimization process. The aim was to select the subset that maximizes the genetic diversity.
- Genome-based models were run using the R-package breedR for each scenario using the simulated data and the accuracy was assessed through a cross-validation routine (the 800 individuals were selected as the T_set_ and the remaining ∼2000 individuals from the trial were in the V_set_). For each sampling scenario we assessed both global (**Appendix Fig.10**) and within-family accuracy (**Appendix Fig.11**).

**Fig.10.**
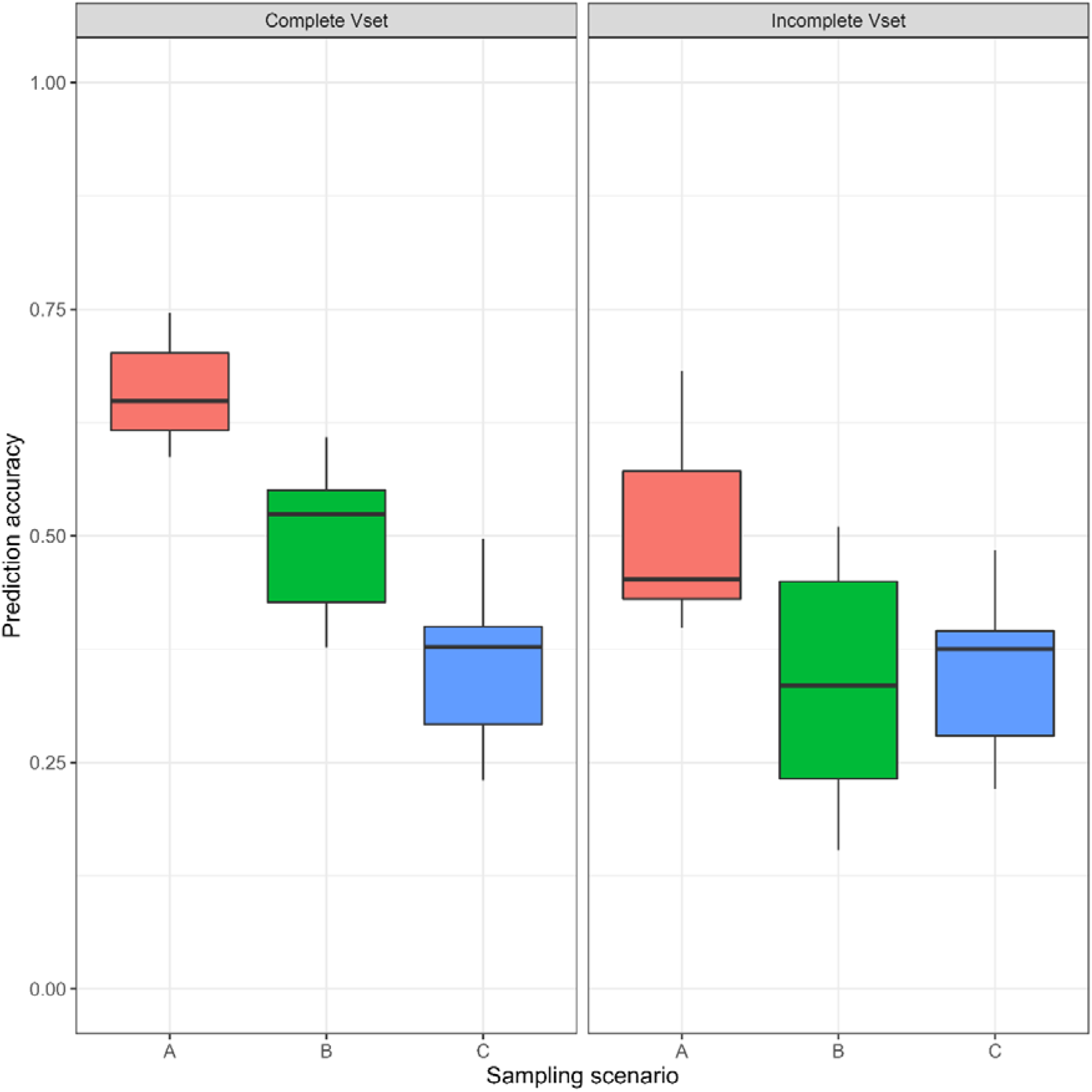
Genome-based prediction accuracy at the global level for the different sampling scenarios. *A: 40 families x 20 individuals, B: 32 families x 25 individuals, C: 20 families x 40 individuals. “Complete Vset” indicates that all individuals in the Vset were considered to assess prediction accuracy, while “incomplete Vset” indicates that only individuals from families not included in the Tset were considered to assess prediction accuracy*.

**Fig.11.**
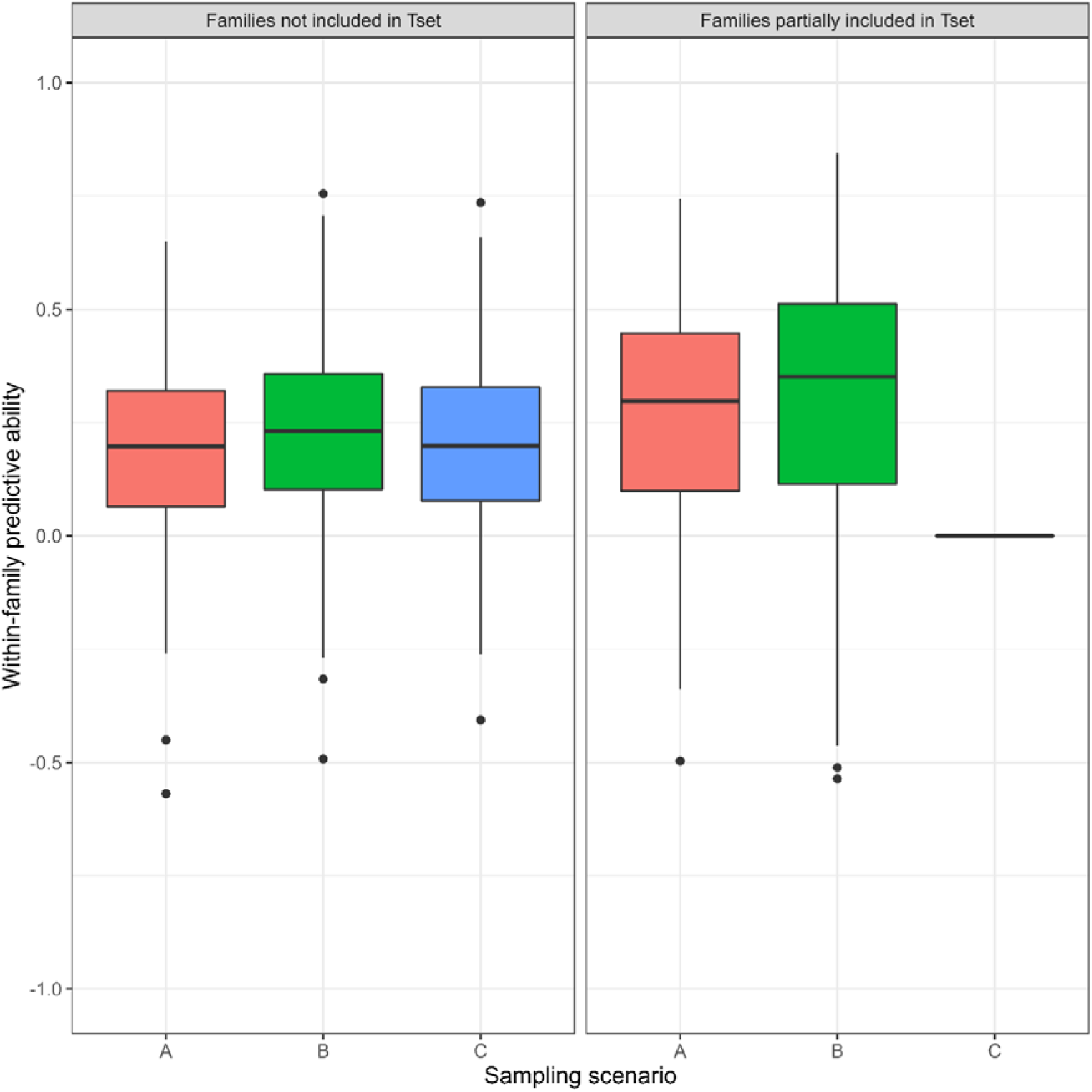
Genome-based within-family prediction accuracy for the different sampling scenarios. *A: 40 families x 20 individuals, B: 32 families x 25 individuals, C: 20 families x 40 individuals. On the left-hand side, assessing within-family accuracy for all the families not included in the Tset, and on the right-hand side only for families with some individuals in the Tset. In the latter modality, prediction accuracy is not available considering sampling scenario C since all the individuals in these families are included in the Tset*.

Finally, the scenario using 40 families with 20 individuals each, was chosen since it achieved significantly higher genome-based prediction accuracy at the global level, and similar within-family accuracy compared to the other 2 scenarios.

### Choice of individuals within each family

Within each of the 40 selected families, we applied the Kennard-Stone algorithm (Kennard and Stone 1969) implemented in the R-package prospectr (Stevens 2022) to get the subset of 20 individuals that maximize phenotypic diversity (now based on the real phenotypic values) for the three traits of interest: circumference, height, and deviation to verticality at 8 years.

### Addition of 200 individuals

Subsequently, additional 200 individuals could be genotyped and added to the study. As the sampling of the 800 individuals was already performed, we chosen to add 20 additional individuals in the 10 out of the 40 selected families. These families (in the main manuscript labeled as the “large families”) were finally made up of 40 individuals each.

These 10 families were chosen to provide a contrast in connectivity within the population sampled. They represent the top 5 and bottom 5 families in terms of connectivity, with their average relatedness to the rest of the families sampled calculated as 0.03 and 0.01, respectively, based on pedigree data. Note that this choice of large families was made a posteriori so that the gap in relatedness with the rest of the population between the two groups of large families may be not very pronounced.

## Appendix 2: simulation of the French maritime pine breeding programme

We simulated a diploid maritime pine genome composed of 12 chromosomes each with a physical length of 2.14e+09bp (Chagné 2002), a genetic length of 1.2M (Chancerel 2013) and a total number of segregating sites of 6910 (10 times the average number of SNP per chromosome available in the real dataset plus 50 theoretical QTL). A simple species-specific demography was used to mimic the selection in the natural environment of the breeding programme base population (popG0): from a wild ancestral population in the Landes forest, we simulated one significant bottleneck reducing effective population size from ∼50000 (Milesi et al., 2023) to 100 (actual effective size for popG0) and with a mutation rate of 4e-18 per generation (Jaramillo-Correa 2020). We investigated the coherence in LD and allele frequency distributions for the 600 individuals of popG0, by comparing real genotyping data and simulated SNP array data (**Appendix Fig.12**). One phenotypic trait was simulated with reference values equal to those of the “height” trait targeted in the breeding programme (phenotypic average=6.81m and genetic variance=0.12m²).

**Fig.12.**
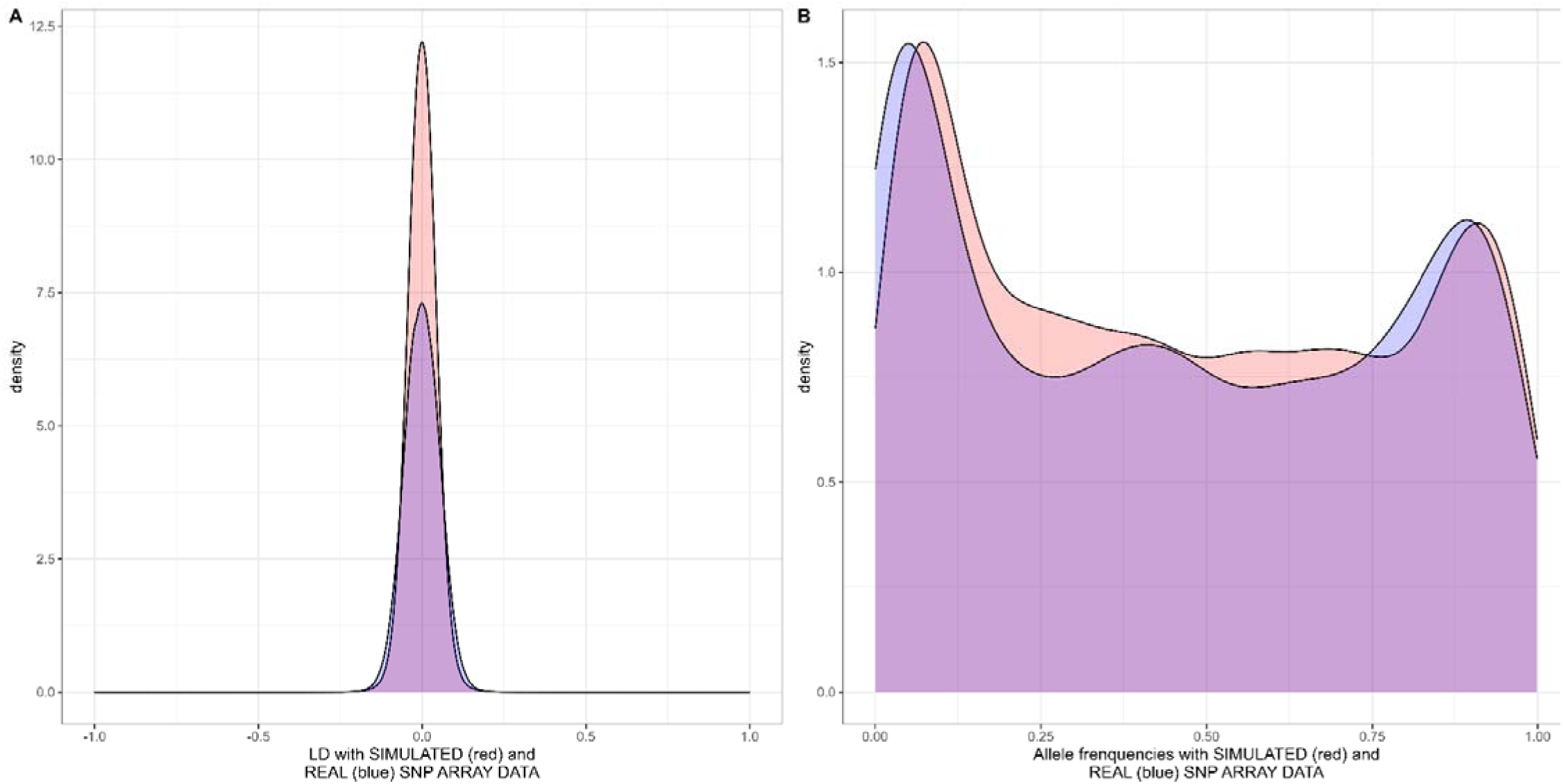
Comparison of LD (A) and allele frequency (B) distributions between real and simulated SNP array data.

The 833 individuals of POP_R_ considered in this study were simulated after the two breeding cycles, as closely as possible to the real-life conditions (**Appendix Fig.13**). Each individual in popG0 took the identity of a real individual in the pedigree, the one with which it shared the same rank in terms of EBV. The EBV were generated with a correlation of 0.97 with true breeding values (BV) since the accuracy of EBV in real programme for these individuals is very high due to progeny testing. Individuals of popG0 were crossed according to the actual crossing plan to generate popP1 FS families of 130 individuals. Individuals for popG1 were selected based on EBV in each family with an intensity of 1.07. The use of this selection intensity, estimated with real data, made it possible to mimic the multi-character aspect of the actual selection carried out, as well as the diversity constraint actually used. Finally, the families of POP_S_ (simulated version of POP_R_) were obtained by crossing the individuals of popG1 according to the actual pedigree.

**Fig.13.**
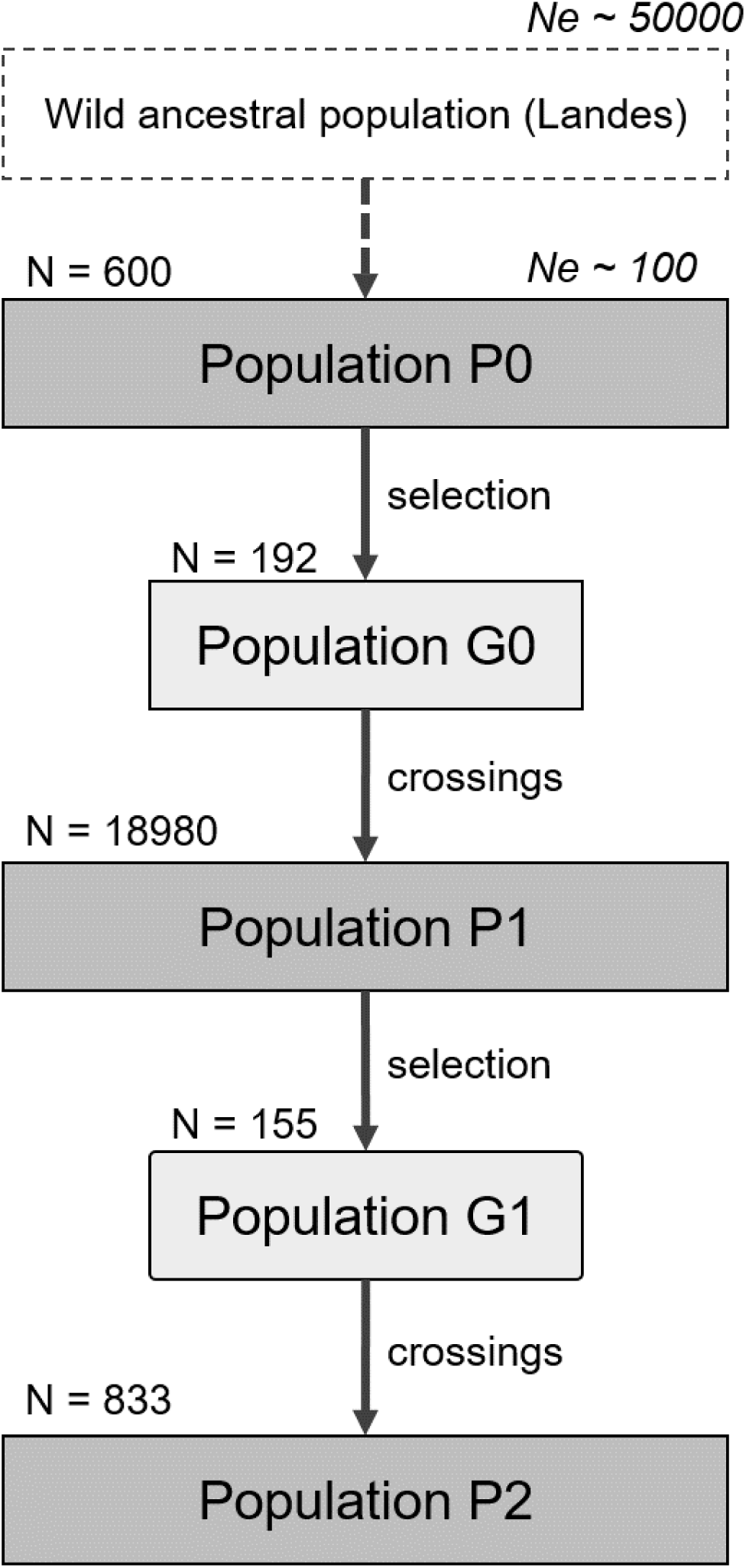
Breeding cycles simulated to mimic the French maritime pine breeding programme.

## Appendix 3: Complementary analysis of genome-based prediction accuracy using deterministic approaches

Mathematical modeling approaches are useful tools to predict the accuracy of genome-based prediction as a function of various population parameters (Daetwyler et al., 2008; Goddard, 2009; Hayes, Visscher, et al., 2009). Deterministic formulas have already been applied to forest trees (Grattapaglia & Resende, 2011) and are here adapted with input parameters from our maritime pine case study, following Gorjanc et al. (2015). The accuracy (square root of reliability) of GEBV was obtained by:

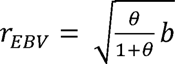

Where *θ* = *nT_set_bh^2^/M_b_* with nT_set_ the size of the training set and h² the heritability of the trait. *M_e_* and *b*, respectively the effective number of chromosome segments and the proportion of genetic variance captured by markers, being defined by *M_e_* = 2*N_e_LC*/log (*N_e_L*) and *b* = *nSNP*/(*nSNP* + *M_e_*), with *N_e_* = 100 the effective size of our breeding population, *L* = 1.2 the average size of maritime pine chromosomes (in Morgans) and *C* = 12 the number of chromosomes. Based on progeny records, the prediction accuracy of non-phenotypes progeny with ABLUP depends solely on the accuracy of EBV in its parents. In case of no selection among parents, we have 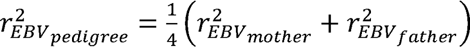 with 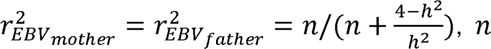 being the number of progenies with phenotypic values per parent.

The trends observed (**Appendix Fig.14**) are fully consistent with those obtained with stochastic simulations presented in the previous section (**Fig.8**). Overall prediction accuracy is mainly impacted by T_set_ size and to a much lesser extent by trait heritability. The advantage of GBLUP over ABLUP models in terms of forward prediction accuracy only becomes apparent above 2000 individuals in the T_set_. Although deterministic approaches are very interesting for quickly revealing key parameters for genome-based prediction accuracy, they have been called into question and refined several times (Brard & Ricard, 2015; Elsen, 2016, 2017). Especially when employing genome-based prediction within full-sibling families, as in this study, accurately forecasting the performance of corresponding accuracy for a specific trait within a particular family remains a challenging endeavor (Schopp et al., 2017)

**Fig.14.**
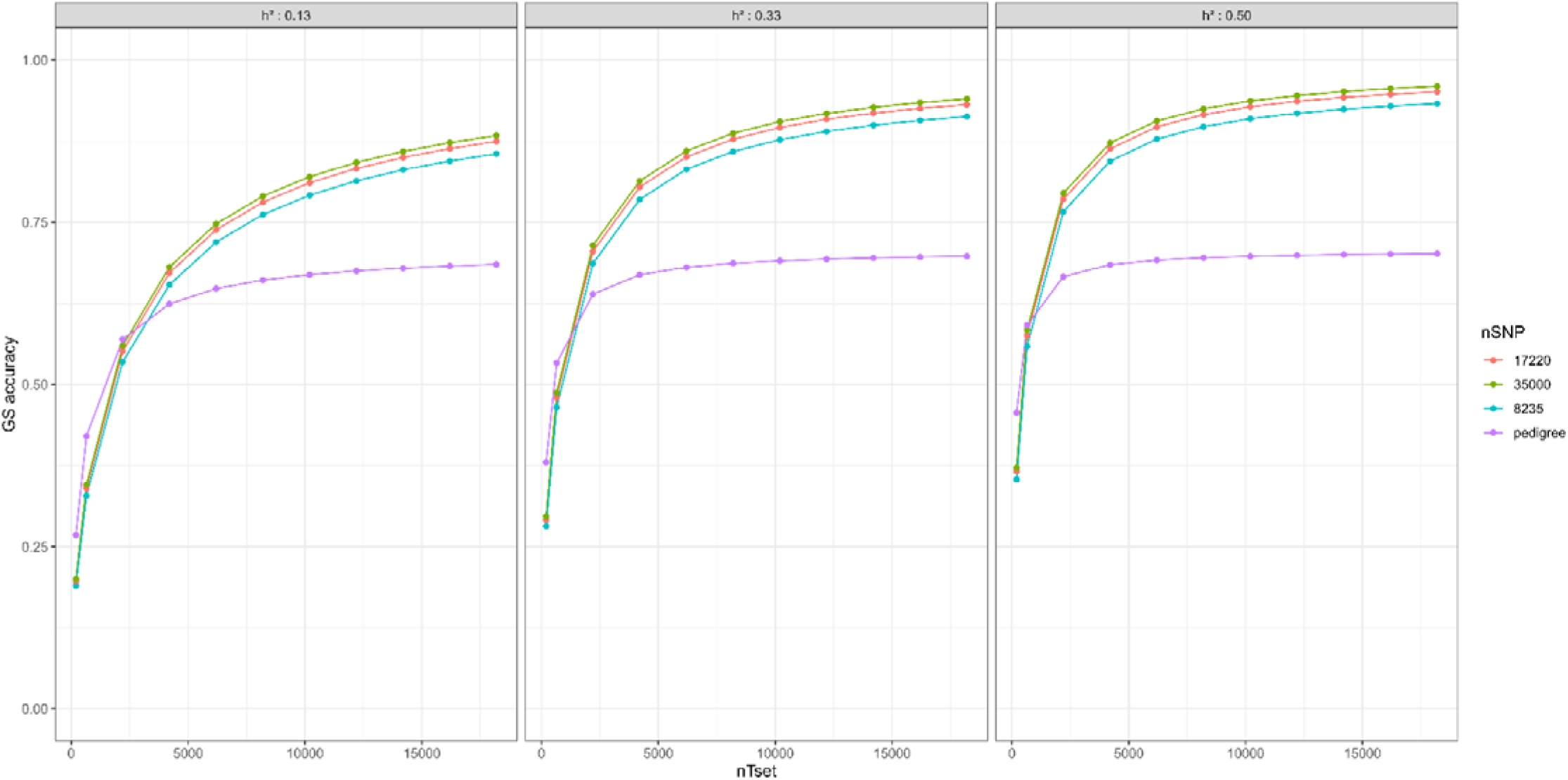
Genome-based accuracy determined with deterministic formulas.

## Declarations

### Ethics approval and consent to participate

Not applicable

### Consent for publication

Not applicable

### Availability of data and material

The datasets generated during and/or analysed during the current study are available in the DATA INARE repository:

https://entrepot.recherche.data.gouv.fr/privateurl.xhtml?token=84e5b92b-eedc-4683-835f-9c7ec44165f4

The custom code and/or software application generated during the current study are available in the Github.com repository, https://github.com/HighlanderLab/vpapin_pine_gs

### Authors’ contributions

**Conceptualization**: Victor Papin, Laurent Bouffier, Leopoldo Sanchez; **Methodology**: Victor Papin, Gregor Gorjanc, Ivan Pocrnic, Laurent Bouffier, Leopoldo Sanchez; **Formal analysis and investigation**: Victor Papin; **Software**: Victor Papin, Gregor Gorjanc, Ivan Pocrnic; **Writing - original draft preparation**: Victor Papin; **Writing - review and editing**: Gregor Gorjanc, Ivan Pocrnic, Laurent Bouffier, Leopoldo Sanchez; **Funding acquisition**: Laurent Bouffier, Leopoldo Sanchez; **Resources**: Laurent Bouffier, Leopoldo Sanchez; **Supervision**: Gregor Gorjanc, Ivan Pocrnic, Laurent Bouffier, Leopoldo Sanchez.

The authors read and approved the final manuscript.

## Acknowledgements

The authors would like to thank GIS “Groupe Pin Maritime du Futur” and INRAE - UEFP (https://doi.org/10.15454/1.5483264699193726E12) for the installation of the studied sites, the management of the sites, the help to collect data (HT and DEV measurements) and biological material (needles). Part of the experiments (DNA extraction, quantification and manipulation) were also performed at the PGTB (doi:10.15454/1.5572396583599417E12), with the help of Mathilde Flores, Christophe Boury and Céline Lalanne. Authors are also thankful to Christophe Plomion for his help during the conceptualization of this study.

## References

Abad Viñas, R., Caudullo, G., Oliveira, S., & de Rigo, D. (2016). Pinus pinaster in Europe□: Distribution, habitat, usage and threats.

Allier, A., Teyssèdre, S., Lehermeier, C., Claustres, B., Maltese, S., Melkior, S., Moreau, L., & Charcosset, A. (2019). Assessment of breeding programs sustainability□: Application of phenotypic and genomic indicators to a North European grain maize program. Theoretical and Applied Genetics, 132(5), 1321□1334. 10.1007/s00122-019-03280-w

Amadeu, R. R., Cellon, C., Olmstead, J. W., Garcia, A. A. F., Resende Jr., M. F. R., & Muñoz, P. R. (2016). AGHmatrix□: R package to construct relationship matrices for autotetraploid and diploid species□: A blueberry example. The Plant Genome, 9(3), 1□10. 10.3835/plantgenome2016.01.0009

Bartholomé, J., Van Heerwaarden, J., Isik, F., Boury, C., Vidal, M., Plomion, C., & Bouffier, L. (2016). Performance of genomic prediction within and across generations in maritime pine. BMC Genomics, 17(1), 604. 10.1186/s12864-016-2879-8

Beaulieu, J., Doerksen, T. K., MacKay, J., Rainville, A., & Bousquet, J. (2014). Genomic selection accuracies within and between environments and small breeding groups in white spruce. BMC Genomics, 15(1), 1048. 10.1186/1471-2164-15-1048

Bernardo, R. (1994). Prediction of maize single-cross performance using RFLPs and information from related hybrids. Crop Science, 34, 20–25. 10.2135/cropsci1994.0011183X003400010003x

Bouffier, L., Raffin, A. A., & Dutkowski, G. (2016, mars 14). Using pedigree and trait relationships to increase gain in the French maritime pine breeding program. IUFRO Conference « Forest Genetics for Productivity ». https://hal.inrae.fr/hal-02801580

Brard, S., & Ricard, A. (2015). Is the use of formulae a reliable way to predict the accuracy of genomic selection? Journal of Animal Breeding and Genetics = Zeitschrift Fur Tierzuchtung Und Zuchtungsbiologie, 132(3), 207□217. 10.1111/jbg.12123

Chagné, D., Lalanne, C., Madur, D., Kumar, S., Frigério, J.-M., Krier, C., Decroocq, S., Savouré, A., Bou-Dagher-Kharrat, M., Bertocchi, E., Brach, J., & Plomion, C. (2002). A high density genetic map of maritime pine based on AFLPs. Annals of Forest Science, 59(5□6), 627□636. 10.1051/forest:2002048

Chancerel, E., Lamy, J.-B., Lesur, I., Noirot, C., Klopp, C., Ehrenmann, F., Boury, C., Provost, G. L., Label, P., Lalanne, C., Léger, V., Salin, F., Gion, J.-M., & Plomion, C. (2013). High-density linkage mapping in a pine tree reveals a genomic region associated with inbreeding depression and provides clues to the extent and distribution of meiotic recombination. BMC Biology, 11(1), 50. 10.1186/1741-7007-11-50

Chen, Z.-Q., Baison, J., Pan, J., Karlsson, B., Andersson, B., Westin, J., García-Gil, M. R., & Wu, H. X. (2018). Accuracy of genomic selection for growth and wood quality traits in two control-pollinated progeny trials using exome capture as the genotyping platform in Norway spruce. BMC Genomics, 19(1), 946. 10.1186/s12864-018-5256-y

Cros, D., Mbo-Nkoulou, L., Bell, J. M., Oum, J., Masson, A., Soumahoro, M., Tran, D. M., Achour, Z., Le Guen, V., & Clement-Demange, A. (2019). Within-family genomic selection in rubber tree (Hevea brasiliensis) increases genetic gain for rubber production. Industrial Crops and Products, 138, 111464. 10.1016/j.indcrop.2019.111464

Crossa, J., Pérez-Rodríguez, P., Cuevas, J., Montesinos-López, O., Jarquín, D., Campos, G. de los, Burgueño, J., González-Camacho, J. M., Pérez-Elizalde, S., Beyene, Y., Dreisigacker, S., Singh, R., Zhang, X., Gowda, M., Roorkiwal, M., Rutkoski, J., & Varshney, R. K. (2017). Genomic Selection in Plant Breeding□: Methods, Models, and Perspectives. Trends in Plant Science, 22(11), 961□975. 10.1016/j.tplants.2017.08.011

Daetwyler, H. D., Villanueva, B., & Woolliams, J. A. (2008). Accuracy of Predicting the Genetic Risk of Disease Using a Genome-Wide Approach. PLOS ONE, 3(10), e3395. 10.1371/journal.pone.0003395

Dehli Vigeland, M. (2022). pedtools□: Creating and Working with Pedigrees and Marker Data (R package version 1.3.0) [Logiciel]. https://github.com/magnusdv/pedtools

Doerksen, T. K., & Herbinger, C. M. (2010). Impact of reconstructed pedigrees on progeny-test breeding values in red spruce. Tree Genetics & Genomes, 6(4), 591□600. 10.1007/s11295-010-0274-1

Durán, R., Isik, F., Zapata-Valenzuela, J., Balocchi, C., & Valenzuela, S. (2017). Genomic predictions of breeding values in a cloned Eucalyptus globulus population in Chile. Tree Genetics & Genomes, 13(4), 74. 10.1007/s11295-017-1158-4

Durel, C.-E. (1992). Gains génétiques attendus après sélection sur index en seconde génération d’amélioration du Pin maritime. Revue forestière française, 44(4), 341□355. 10.4267/2042/26331

Eckert, A. J., van Heerwaarden, J., Wegrzyn, J. L., Nelson, C. D., Ross-Ibarra, J., González-Martínez, S. C., & Neale, D. B. (2010). Patterns of Population Structure and Environmental Associations to Aridity Across the Range of Loblolly Pine (Pinus taeda L., Pinaceae). Genetics, 185(3), 969□982. 10.1534/genetics.110.115543

El-Dien, O. G., Ratcliffe, B., Klápště, J., Porth, I., Chen, C., & El-Kassaby, Y. A. (2018). Multienvironment genomic variance decomposition analysis of open-pollinated Interior spruce (Picea glauca x engelmannii). Molecular Breeding, 38(3), 26. 10.1007/s11032-018-0784-3

Elsen, J.-M. (2016). Approximated prediction of genomic selection accuracy when reference and candidate populations are related. Genetics Selection Evolution, 48(1), 18. 10.1186/s12711-016-0183-3

Elsen, J.-M. (2017). An analytical framework to derive the expected precision of genomic selection. Genetics Selection Evolution, 49(1), 95. 10.1186/s12711-017-0366-6

Fuentes-Utrilla, P., Goswami, C., Cottrell, J. E., Pong-Wong, R., Law, A., A’Hara, S. W., Lee, S. J., & Woolliams, J. A. (2017). QTL analysis and genomic selection using RADseq derived markers in Sitka spruce□: The potential utility of within family data. Tree Genetics & Genomes, 13(2), 33. 10.1007/s11295-017-1118-z

Gaynor, R. C., Gorjanc, G., & Hickey, J. M. (2021). AlphaSimR□: An R package for breeding program simulations. G3 Genes|Genomes|Genetics, 11(2), jkaa017. 10.1093/g3journal/jkaa017

Goddard, M. (2009). Genomic selection□: Prediction of accuracy and maximisation of long term response. Genetica, 136(2), 245□257. 10.1007/s10709-008-9308-0

Gorjanc, G., Bijma, P., & Hickey, J. M. (2015). Reliability of pedigree-based and genomic evaluations in selected populations. Genetics Selection Evolution, 47(1), 65. 10.1186/s12711-015-0145-1

Gorjanc, G., Gaynor, R.C., & Hickey, J.M. (2018). Optimal cross selection for long-term genetic gain in two-part programs with rapid recurrent genomic selection. Theor Appl Genet, 131, 1953– 1966. 10.1007/s00122-018-3125-3

Grattapaglia, D., & Resende, M. D. V. (2011). Genomic selection in forest tree breeding. Tree Genetics & Genomes, 7(2), 241□255. 10.1007/s11295-010-0328-4

Grattapaglia, D., Vilela Resende, M. D., Resende, M. R., Sansaloni, C. P., Petroli, C. D., Missiaggia, A. A., Takahashi, E. K., Zamprogno, K. C., & Kilian, A. (2011). Genomic Selection for growth traits in Eucalyptus□: Accuracy within and across breeding populations. BMC Proceedings, 5(7), O16. 10.1186/1753-6561-5-S7-O16

Guilbaud, R., Biselli, C., Buiteveld, J., Cattivelli, L., Copini, P., Dowkiw, A., Esselink, D., Fricano, A., Guerin, V., Jorge, V., & others. (2020). Development of a new tool (4TREE) for adapted genome selection in European tree species. Proceedings of the Gentree Symposium. Proceedings of the Gentree Symposium, Avignon, France.

Habier, D., Fernando, R. L., & Dekkers, J. C. M. (2007). The Impact of Genetic Relationship Information on Genome-Assisted Breeding Values. Genetics, 177(4), 2389□2397. 10.1534/genetics.107.081190

Habier, D., Fernando, R. L., & Garrick, D. J. (2013). Genomic BLUP Decoded□: A Look into the Black Box of Genomic Prediction. Genetics, 194(3), 597□607. 10.1534/genetics.113.152207

Hallander, J., & Waldmann, P. (2009). Optimum contribution selection in large general tree breeding populations with an application to Scots pine. Theoretical and Applied Genetics, 118(6), 1133□1142. 10.1007/s00122-009-0968-7

Hayes, B. J., Bowman, P. J., Chamberlain, A. J., & Goddard, M. E. (2009). Invited review□: Genomic selection in dairy cattle: Progress and challenges. Journal of Dairy Science, 92(2), 433□443. 10.3168/jds.2008-1646

Hayes, B. J., Visscher, P. M., & Goddard, M. E. (2009). Increased accuracy of artificial selection by using the realized relationship matrix. Genetics Research, 91(1), 47□60. 10.1017/S0016672308009981

Isik, F., Bartholomé, J., Farjat, A., Chancerel, E., Raffin, A., Sanchez, L., Plomion, C., & Bouffier, L. (2016). Genomic selection in maritime pine. Plant Science, 242, 108□119. 10.1016/j.plantsci.2015.08.006

Iwata, H., Hayashi, T., & Tsumura, Y. (2011). Prospects for genomic selection in conifer breeding□: A simulation study of Cryptomeria japonica. Tree Genetics & Genomes, 7(4), 747□758. 10.1007/s11295-011-0371-9

Jannink, J.-L. (2010). Dynamics of long-term genomic selection. Genetics Selection Evolution, 42(1), 35. 10.1186/1297-9686-42-35

Jaramillo-Correa, J. P., Bagnoli, F., Grivet, D., Fady, B., Aravanopoulos, F. A., Vendramin, G. G., & González-Martínez, S. C. (2020). Evolutionary rate and genetic load in an emblematic Mediterranean tree following an ancient and prolonged population collapse. Molecular Ecology, 29(24), 4797□4811. 10.1111/mec.15684

Karaman, E., Cheng, H., Firat, M. Z., Garrick, D. J., & Fernando, R. L. (2016). An Upper Bound for Accuracy of Prediction Using GBLUP. PLOS ONE, 11(8), e0161054. 10.1371/journal.pone.0161054

Kujala, S. T., & Savolainen, O. (2012). Sequence variation patterns along a latitudinal cline in Scots pine (Pinus sylvestris)□: Signs of clinal adaptation? Tree Genetics & Genomes, 8(6), 1451□1467. 10.1007/s11295-012-0532-5

Lebedev, V. G., Lebedeva, T. N., Chernodubov, A. I., & Shestibratov, K. A. (2020). Genomic selection for forest tree improvement□: Methods, achievements and perspectives. Forests, 11(11), 1190. 10.3390/f11111190

Legarra, A., Robert-Granié, C., Manfredi, E., & Elsen, J.-M. (2008). Performance of Genomic Selection in Mice. Genetics, 180(1), 611□618. 10.1534/genetics.108.088575

Lenz, P. R. N., Beaulieu, J., Mansfield, S. D., Clément, S., Desponts, M., & Bousquet, J. (2017). Factors affecting the accuracy of genomic selection for growth and wood quality traits in an advanced-breeding population of black spruce (Picea mariana). BMC Genomics, 18(1), 335. 10.1186/s12864-017-3715-5

Lenz, P. R. N., Nadeau, S., Azaiez, A., Gérardi, S., Deslauriers, M., Perron, M., Isabel, N., Beaulieu, J., & Bousquet, J. (2020). Genomic prediction for hastening and improving efficiency of forward selection in conifer polycross mating designs□: An example from white spruce. Heredity, 124(4), Article 4. 10.1038/s41437-019-0290-3

Lenz, P. R. N., Nadeau, S., Mottet, M.-J., Perron, M., Isabel, N., Beaulieu, J., & Bousquet, J. (2020). Multi-trait genomic selection for weevil resistance, growth, and wood quality in Norway spruce. Evolutionary Applications, 13(1), 76□94. 10.1111/eva.12823

Li, Y., Klápště, J., Telfer, E., Wilcox, P., Graham, N., Macdonald, L., & Dungey, H. S. (2019). Genomic selection for non-key traits in radiata pine when the documented pedigree is corrected using DNA marker information. BMC Genomics, 20(1), 1026. 10.1186/s12864-019-6420-8

Lindgren, D., Gea, L., & Jefferson, P. A. (1996). Loss of genetic diversity monitored by status number. Silvae Genetica, 45, 52□59.

Meuwissen, T. H. E. (1997). Maximizing the response of selection with a predefined rate of inbreeding. Journal of Animal Science, 75(4), 934□940. 10.2527/1997.754934x

Meuwissen, T. H. E., Hayes, B. J., & Goddard, M. E. (2001). Prediction of Total Genetic Value Using Genome-Wide Dense Marker Maps. Genetics, 157(4), 1819□1829. 10.1093/genetics/157.4.1819

Milesi, P., Kastally, C., Dauphin, B., Cervantes, S., Bagnoli, F., Budde, K. B., Cavers, S., Fady, B., Faivre-Rampant, P., & Gonzalez-Martinez, S. C. (2023). Synchronous effective population size changes and genetic stability of forest trees through glacial cycles. BioRxiv. [Preprint]. 10.1101/2023.01.05.522822. Accessed January 2023.

Mrode, R. A., & Pocrnic, I. (2023). Linear models for the prediction of the genetic merit of animals. 4^th^ edition, United Kingdom:CABI, 412p. 10.1079/9781800620506.0000

Muñoz, F., & Sanchez, L. (2020). breedR: statistical methods for forest genetic resources analysts. R package Version 0.12-5. https://github.com/famuvie/breedR

Munoz, P. R., Resende Jr., M. F. R., Huber, D. A., Quesada, T., Resende, M. D. V., Neale, D. B., Wegrzyn, J. L., Kirst, M., & Peter, G. F. (2014). Genomic Relationship Matrix for Correcting Pedigree Errors in Breeding Populations□: Impact on Genetic Parameters and Genomic Selection Accuracy. Crop Science, 54(3), 1115□1123. 10.2135/cropsci2012.12.0673

Nejati-Javaremi, A., Smith, C., & Gibson, J. P. (1997). Effect of total allelic relationship on accuracy of evaluation and response to selection. Journal of Animal Science, 75(7), 1738–1745. 10.2527/1997.7571738x

Pégard, M., Segura, V., Muñoz, F., Bastien, C., Jorge, V., & Sanchez, L. (2020). Favorable Conditions for Genomic Evaluation to Outperform Classical Pedigree Evaluation Highlighted by a Proof-of-Concept Study in Poplar. Frontiers in Plant Science, 11. https://www.frontiersin.org/articles/10.3389/fpls.2020.581954

Plomion, C., Chancerel, E., Endelman, J., Lamy, J.-B., Mandrou, E., Lesur, I., Ehrenmann, F., Isik, F., Bink, M. C., van heerwaarden, J., & Bouffier, L. (2014). Genome-wide distribution of genetic diversity and linkage disequilibrium in a mass-selected population of maritime pine. BMC Genomics, 15(1), 171. 10.1186/1471-2164-15-171

Pryce, J. E., Daetwyler, H. D., Pryce, J. E., & Daetwyler, H. D. (2011). Designing dairy cattle breeding schemes under genomic selection□: A review of international research. Animal Production Science, 52(3), 107□114. 10.1071/AN11098

R Core Team. (2022). R: A Language and Environment for Statistical Computing [Logiciel]. https://www.R-project.org/

Ratcliffe, B., El-Dien, O. G., Klápště, J., Porth, I., Chen, C., Jaquish, B., & El-Kassaby, Y. A. (2015). A comparison of genomic selection models across time in interior spruce (Picea engelmannii × glauca) using unordered SNP imputation methods. Heredity, 115(6), 547□555. 10.1038/hdy.2015.57

Rauf, S., da Silva, J. T., Khan, A. A., & Naveed, A. (2010). Consequences of plant breeding on genetic diversity. International Journal of plant breeding, 4(1), 1□21.

Resende, J., M. F. R., Muñoz, P., Resende, M. D. V., Garrick, D. J., Fernando, R. L., Davis, J. M., Jokela, E. J., Martin, T. A., Peter, G. F., & Kirst, M. (2012). Accuracy of Genomic Selection Methods in a Standard Data Set of Loblolly Pine (Pinus taeda L.). Genetics, 190(4), 1503□1510. 10.1534/genetics.111.137026

Resende Jr, M. F. R., Muñoz, P., Acosta, J. J., Peter, G. F., Davis, J. M., Grattapaglia, D., Resende, M. D. V., & Kirst, M. (2012). Accelerating the domestication of trees using genomic selection□: Accuracy of prediction models across ages and environments. New Phytologist, 193(3), 617□624. 10.1111/j.1469-8137.2011.03895.x

Resende, R. T., Resende, M. D. V., Silva, F. F., Azevedo, C. F., Takahashi, E. K., Silva-Junior, O. B., & Grattapaglia, D. (2017). Assessing the expected response to genomic selection of individuals and families in Eucalyptus breeding with an additive-dominant model. Heredity, 119(4), Article 4. 10.1038/hdy.2017.37

Schopp, P., Müller, D., Wientjes, Y. C. J., & Melchinger, A. E. (2017). Genomic Prediction Within and Across Biparental Families□: Means and Variances of Prediction Accuracy and Usefulness of Deterministic Equations. G3 Genes|Genomes|Genetics, 7(11), 3571□3586. 10.1534/g3.117.300076

Stranden, I., & Garrick, D. J. (2009). Derivation of equivalent computing algorithms for genomic predictions and reliabilities of animal merit. Journal of Dairy Science, 92(6), 2971–2975. 10.3168/jds.2008-1929

Thistlethwaite, F. R., Ratcliffe, B., Klápště, J., Porth, I., Chen, C., Stoehr, M. U., & El-Kassaby, Y. A. (2017). Genomic prediction accuracies in space and time for height and wood density of Douglas-fir using exome capture as the genotyping platform. BMC Genomics, 18(1), 930. 10.1186/s12864-017-4258-5

Thistlethwaite, F. R., Ratcliffe, B., Klápště, J., Porth, I., Chen, C., Stoehr, M. U., & El-Kassaby, Y. A. (2019). Genomic selection of juvenile height across a single-generational gap in Douglas-fir. Heredity, 122(6), Article 6. 10.1038/s41437-018-0172-0

Ukrainetz, N. K., & Mansfield, S. D. (2019). Assessing the sensitivities of genomic selection for growth and wood quality traits in lodgepole pine using Bayesian models. Tree Genetics & Genomes, 16(1), 14. 10.1007/s11295-019-1404-z

Vidal, M., Plomion, C., Raffin, A., Harvengt, L., & Bouffier, L. (2017). Forward selection in a maritime pine polycross progeny trial using pedigree reconstruction. Annals of Forest Science, 74(1), 21. 10.1007/s13595-016-0596-8

Woolliams, J. a., Berg, P., Dagnachew, B. s., & Meuwissen, T. h. e. (2015). Genetic contributions and their optimization. Journal of Animal Breeding and Genetics, 132(2), 89□99. 10.1111/jbg.12148

Zapata-Valenzuela, J., Isik, F., Maltecca, C., Wegrzyn, J., Neale, D., McKeand, S., & Whetten, R. (2012). SNP markers trace familial linkages in a cloned population of Pinus taeda—Prospects for genomic selection. Tree Genetics & Genomes, 8(6), 1307□1318. 10.1007/s11295-012-0516-5

Zapata-Valenzuela, J., Whetten, R. W., Neale, D., McKeand, S., & Isik, F. (2013). Genomic Estimated Breeding Values Using Genomic Relationship Matrices in a Cloned Population of Loblolly Pine. G3 Genes|Genomes|Genetics, 3(5), 909□916. 10.1534/g3.113.005975

Zhou, L., Chen, Z., Olsson, L., Grahn, T., Karlsson, B., Wu, H. X., Lundqvist, S.-O., & García-Gil, M. R. (2020). Effect of number of annual rings and tree ages on genomic predictive ability for solid wood properties of Norway spruce. BMC Genomics, 21(1), 323. 10.1186/s12864-020-6737-3

